# Emotions and bodily sensations evoked by cardiac adenosine stress

**DOI:** 10.64898/2026.06.15.731610

**Authors:** Ksenia Egorova, Teemu Maaniitty, Henri Kärpijoki, Antti Saraste, Juhani Knuuti, Lauri Nummenmaa

## Abstract

Peripheral physiological processes contribute to emotions; however, the extent of the influence of internal signals on subjective emotional experiences remains a subject of debate.

**Methods:** Here we investigated the bodily sensations induced by pharmacological stimulation with adenosine, a potent vasodilator that does not cross the blood-brain barrier, and how these sensations are translated into emotional experiences in 193 patients (94 males, 99 females; mean age 62.6 ± 9.2 years) undergoing pharmacological cardiac stress testing as a part of their clinical assessment for suspected obstructive coronary artery disease. We characterized the subjective experiences induced by adenosine using topographical bodily maps and Likert-rating scales of 16 somatic sensations and 16 emotions.

**Results:** Adenosine induced strong and consistent bodily sensations related to dyspnea, chest pain, tachycardia, feelings of warmth, and tension in the body. The induced sensations were topographically distinct, with pain and pressure concentrated in the chest, warmth distributed across the upper body, and weakness localized to the torso. These sensations were accompanied by a significant increase in negative emotions, such as anxiety, fear, restlessness, panic, and a decrease in positive emotions of happiness, joy, and calmness. Peak heart rate response predicted the intensity of most bodily and emotional responses. Patients diagnosed with myocardial ischemia did not show heightened sensitivity to adenosine-induced bodily sensations compared to non-ischemic patients, despite exhibiting a blunted tachycardiac reflex.

**Conclusion:** These findings show that peripheral physiological activation alone is sufficient to drive subjective emotional experiences, providing causal evidence for the embodied nature of emotions.

## Introduction

Emotions coordinate behaviour and physiological states during survival-salient events by adjusting skeletomuscular, neuroendocrine, autonomic, and central nervous system activity for maximizing survival needs (Damasio & Carvalho, 2013; Nummenmaa & Saarimäki, 2019). The physiological background can shift our affective state and behavioural priorities, adjusting the perception of what is currently salient (Critchley & Harrison, 2013). This coordinated physiological activation pattern is often accompanied by a subjective experience of emotion (such as “I feel afraid”). This subjective component of emotion is shaped by the ongoing physiological state and bodily processes (Barrett & Lindquist, 2008; Critchley & Harrison, 2013), and recent work based on topographical self-reports has established that different emotions are accompanied by distinct “feeling fingerprints” or subjective experience patterns in the body (Nummenmaa et al., 2014, 2018; Putkinen et al., 2024). Thus, the subjective emotional experience presumably arises at least partially through interoception of emotion-related peripheral physiological changes.

The exact contribution of autonomic activity and the corresponding interoceptive signals to emotions and their regulation remains, however, poorly understood, in part due to the lack of experimental models that allow specific manipulation of peripheral interoceptive signals. A major challenge involves isolating the effects of interoceptive signals from concurrent central neural activation and other confounds, and triggering peripheral autonomic activation without directly altering the central nervous system state (Cardenas et al., 2025; Critchley & Harrison, 2013). Recent studies have addressed direct cardiac or sympathetic stimulation and its effects on affect and behavior in animals. Pharmacologic stimulation of the sympathetic nervous system can decrease tolerance to aversive stimuli in a rewarding task (Cardenas et al., 2025) while optogenetic pacing of heart rate can increase anxious behavior of rats in stressful environments (Hsueh et al., 2023). Manipulating interoceptive signals is thus a viable model to study the contribution of autonomic activity and its interoceptive processing on emotion.

Clinical studies indicate a tight linkage between interoception and emotion networks in the brain. Patients with cardiovascular disease show altered brain activity in regions related to pain perception (Wittbrodt et al., 2020), emotional regulation, and working memory during mental and emotional stress tasks (Bremner et al., 2018; Moazzami et al., 2020; Soufer et al., 2009; Tawakol et al., 2017; Wittbrodt et al., 2020), as well as in regions mediating the cardioception and cardiovascular arousal during interoceptive tasks (Yoris et al., 2018). Alterations in cardioception potentially reflect altered sensitivity to cardiac signals and are consistent with predisposition to anxiety and heightened stress responses that are prevalent in cardiovascular patients (Abohashem et al., 2026; Domschke et al., 2010; Korkmaz et al., 2017; Todaro et al., 2007). However, it remains unknown to what extent the physiological changes that accompany cardiovascular disease alter the affective integration and regulation in cardiovascular patients. Understanding this connection is clinically relevant since anxiety and depression are prevalent in cardiovascular patients and known risk factors for worse disease prognosis and major adverse cardiac events (Batelaan et al., 2016; Jiang et al., 2004).

Here, we measured the topographical bodily sensations as well as subjective emotional experiences during vasodilatory pharmacological stress. Adenosine is a potent vasodilator that is routinely used in pharmacological cardiac stress testing to assess myocardial stress perfusion and detect myocardial ischemia. It induces angina-like bodily sensations and dyspnea without crossing the blood-brain barrier or directly stimulating the sympathetic nervous system (Burki et al., 2005; Isakovic et al., 2004). These sensations are produced by vasodilation and stimulation of adenosine receptors in endothelial cells lining the blood vessels, the aortic body (Biaggioni et al., 1987; Reiss et al., 2019), and on the vagal C-fibres in the lungs (Burki & Lee, 2010). The half-life of adenosine in the body is very short, and the physiological effects recover within minutes after bolus administration (Biaggioni et al., 1987). This makes adenosine a well-suited model for investigating how peripheral physiological changes give rise to subjective emotional states in humans.

By comparing individuals with confirmed stress-induced myocardial ischemia to those without, we investigate how the extent of cardiovascular pathology alters the integration of interoceptive signals into emotions. We hypothesized that i. adenosine-related autonomic changes cause topographically specific somatic sensations as well as subjective emotions, and ii. that patients with myocardial ischemia will show heightened responses to uncomfortable and angina-like bodily sensations as compared to patients without ischemia.

## Methods

### Participants

We prospectively recruited 223 patients undergoing clinically indicated myocardial perfusion imaging with pharmacological cardiac stress test (pCST) for evaluation of suspected obstructive coronary artery disease. Most patients had undergone prior coronary computed tomography (CT) angiography showing a coronary artery stenosis with unclear functional significance, and therefore, required functional evaluation by myocardial perfusion imaging, in accordance with clinical guidelines (Vrints et al., 2024). We included the data from participants who completed the pCST scan and all the questionnaires. Eleven participants were excluded due to interrupted, incomplete, or inconclusive pCST, and 19 were excluded due to a history of cerebral stroke, leaving **193 participants** (males = 94, females = 99) with a mean ± SD age of 62.6 ± 9.2 years.

Our sample contained an equivalent number of female and male patients; however, myocardial perfusion imaging revealed that ischemic patients were mostly men (74%). After adjusting for age, the probability of myocardial ischemia was more than 7 times higher in men than in women (OR = 7.57, 95% CI [4.0028–14.88], p < 0.001). Demographic information is described in *Table 1*.

**Table 1.** Demographic information on patients. Data is shown as mean ± SD.

|  | All patients | Ischemic | Non-ischemic |
| --- | --- | --- | --- |
| **Total number** | 193 | 85 | 108 |
| Women N + (%) | 97 (50%) | 22 (26%) | 77 (71%) |
| Age (years) | 62.5 ± 9.2 | 63.0 ± 8.7 | 62.2 ± 9.6 |
| **Cardiovascular factors** |  |  |  |
| CVD medication (N) | 154 | 79 | 78 |
| Rest diastolic pressure (mmHg) | 81.4 ± 12.3 | 79.1 ± 14.3 | 83.2 ± 10.2 |
| Rest systolic pressure (mmHg) | 144.2 ± 22.3 | 142.6 ± 25.1 | 145.4 ± 19.8 |
| **Other health aspects** |  |  |  |
| Body mass index | 30.3 ± 5.3 | 31.3 ± 5.7 | 29.5 ± 4.8 |
| CNS medication (N) | 50 | 24 | 26 |

Information on the medication was collected based on patient interviews and electronic medical records. Cardiovascular disease (CVD) medication mentioned in Table 1 included angiotensin-converting-enzyme inhibitors, angiotensin II receptor blockers, anticoagulants, antiplatelet therapy, antiarrhythmics, beta-blockers, calcium channel blockers, diuretics, long-acting nitrates, and lipid-lowering therapy. Central nervous system (CNS) affecting medications included antidepressants, antipsychotics, hypnotics, benzodiazepines, opioid analgesics, local anesthetics, hydroxyzine, and donepezil.

The Ethics Committee of the Hospital District of Southwest Finland approved the study protocol, which was registered as a clinical trial (Knuuti, 2023) and was completed in accordance with the Declaration of Helsinki at the Turku PET Centre. All patients signed written informed consent.

### Pharmacological stress testing protocol

Participants underwent myocardial perfusion imaging with 15O-labeled water (350 MBq) at rest and with pharmacological stress in a long axial field-of-view (106 cm) PET-CT scanner (Siemens Biograph Vision Quadra) as a part of their clinical assessment. The hybrid positron emission tomography (PET)/CT procedure for detecting myocardial ischemia is performed routinely in the Turku PET Centre and has been described and validated in previous publications (Kajander et al., 2010; Maaniitty et al., 2017). Myocardial ischemia was diagnosed by a supervising and reporting physician, based on previously established cut-off values (Danad et al., 2017).

The imaging procedure consisted of a 4-minute 40-second rest scan followed by a 4-minute 40-second stress scan with pCST using adenosine infusion (0.140 mg/kg/min). The break between rest and stress scans was 10 minutes. Electrocardiography was monitored throughout the scans. Heart rate and brachial blood pressure were collected at resting baseline before the scan and after 3 and 6 minutes of adenosine stress using hospital monitoring equipment (Tango M2, SunTech Medical). The software (CardioSoft Client) automatically extracted the highest observed (peak) heart rate during stress. After the stress PET/CT scan, participants reported the bodily sensations and emotions they experienced during the scans and filled in the personality questionnaires on a tablet computer on the gorilla.sc platform.

### Self-report measures

After PET/CT scans were completed, patients evaluated their emotional state and bodily sensations during the scans using a self-report questionnaire with 16 different emotions (such as anxiety, panic, confusion, stress, joy) and 16 different somatic sensations (such as quickened breathing, pressure in the chest, feeling unwell, headache, muscle tensing). See Supplementary materials (**Appendix 3**) for a complete list of measured emotions and sensations. Participants rated the items on a 0–10-point Likert scale. We also measured the topography of four bodily sensations using the emBODY self-report tool after the scans (Nummenmaa et al., 2014, 2018). In this task, the participants were shown one sensation and a blank silhouette of a human body at a time and asked to color the regions of the body where they experienced the sensation during the scan. Patients reported sensation maps for warmth, pressure, pain, and weakness, which have been previously reported as common side effects of adenosine administration (Cerqueira et al., 1994).

### Analysis

#### Blood pressure and heart rate

We compared the values between ischemic and non-ischemic patients and men and women by fitting a linear mixed model with ischemic status and sex as fixed effects and subjects as random effects.

#### Topographical maps of bodily sensations

We analyzed topographical sensation maps (all *N* = 181) as described previously (Nummenmaa et al., 2014, 2018). Bodily sensation maps were first screened for anomalous response patterns (scribbling, etc.), and responses outside the body area were masked out. To assess where patients felt different sensations (warmth, pressure, pain, and weakness) in the body at resting baseline and adenosine infusion, we used mass univariate t-tests to compare pixelwise activations against zero for each sensation separately. This resulted in four statistical summary maps (t-maps) where pixel intensities reflect the strength of statistically significant convergence of reported sensations across subjects. Averaging across all four maps, we also created a bodily map of cumulative sensations induced by adenosine. We performed pixelwise repeated measure t-tests to reveal regions where the sensations were stronger for the stress versus rest scan. We controlled the false positive rate using the false discovery rate (FDR) (Benjamini & Hochberg, 1995).

#### Bodily sensations elicited by adenosine and associated emotional responses

For the analysis of emotions and bodily sensations (all *N* = 182) evoked by adenosine, we computed change scores (adenosine minus baseline). We conducted a non-parametric Wilcoxon signed-rank test to test the statistical significance of the change in bodily sensations induced by adenosine and the emotional response to them. We employed non-parametric statistical tests due to non-normal data distribution. We tested the effects of myocardial ischemia on self-reports using the Mann-Whitney U-test. We adjusted all p-values for multiple comparisons using the FDR approach (Benjamini & Hochberg, 1995). We analyzed the joint effects of sex and ischemia status using two-way aligned rank-transformed ANOVA, since the variance between the groups was not equal, and the normality of distribution was violated. We set ischemic status (ischemic vs. non-ischemic) and sex as between-subject factors.

#### Associating topographically mapped sensations with a change in the strength of emotional and bodily responses

We next examined whether reported 16 somatic and 16 emotional responses to adenosine were associated with the net reported sensation in the body in each of the four bodily maps. We first computed the total number of pixels colored in the adenosine and rest conditions to index the total experienced sensations and then computed the change scores (adenosine minus baseline). We then used Spearman’s rank correlation analysis to correlate somatic and emotional responses with the net sensation changes in each bodily map. In a complementary methodological approach, we also predicted the pixel-wise sensation changes in the body maps with each of the 16 somatic sensations and 16 different emotions. A detailed description of methods and results can be found in the Supplementary Materials (**Appendix 2**).

#### Associating peak heart rate response with changes in the strength of emotional and bodily responses

To investigate the relationship between peak heart rate response and change in self-reported emotions and sensations, we fitted separate multiple linear regression models for each emotion and bodily sensation. We defined the change in the emotion or sensation rating as the dependent variable and maximum change in the heart rate (peak minus baseline), ischemic status, and sex as predictors. We centered the maximum change in heart rate on the mean and scaled it in 10 bpm units for simpler interpretation of the regression coefficients. We extracted unstandardized regression coefficients (b), computed 95% confidence intervals (CI), and corrected p-values for multiple comparisons (Benjamini & Hochberg, 1995). The same model was applied to HR, and diastolic and systolic blood pressure ratings at the three-minute mark; these results are not reported in the main text, since the three-minute mark measurement is already past the adenosine peak effect (see Supplementary materials Appendix 6.4) (Biaggioni et al., 1987). We performed all analyses in R.

## Results

### Cardiovascular responses to adenosine

Adenosine increased the heart rate in both ischemic and non-ischemic subjects (**Appendix 5**). Mean heart rate increased from the baseline but was not statistically different between three- and six-minute timepoints. Linear mixed effect model revealed a significant effect of sex on the heart rates (*t*(266) = - 3.069, *b*_men_= -8.64, *p*_FDR_= 0.0061), and significant contribution of ischemic status on heart rates at three (*t*(266) = -4.62, *b*_ischemic_= -9.096, *p*_FDR_= 0.0001) and six minutes (*t*(266) = -4.40, b_ischemic_= -8.68, *p*_FDR_ < 0.0001). At three minutes, for ischemic patients the mean heart rate was 79.5 bpm, 95% CI [75.6 - 83.3], which was 13.3 ± 9.1 bpm increase from rest. For patients without stress-induced myocardial ischemia, the mean heart rate was 87.7 bpm, 95% CI [84.3 - 91.9], which was 20.8 ± 11.1 bpm increase from rest. Female patients exhibited higher heart rates at rest and during stress than males. However, the magnitude of adenosine-induced increase in heart rate from rest was not statistically different across sexes.

Both sex (*b*_men_= -7.38, *p-value =* 0.011) and ischemic status (*b*_ischemic_ = -7.86, *p-value* = 0.0070) contribute to the peak heart rate induced by adenosine. The mean sex-adjusted peak heart rate induced by adenosine was 89.1 ± 14.0 bpm for ischemic and 94.2 ± 13.1 bpm for non-ischemic patients.

Linear mixed modelling did not reveal significant effects of sex and ischemic status on the blood pressure responses. Both systolic and diastolic pressures decreased slightly during the stress scan, with the lowest point at the six-minute mark. See detailed results in the Supplementary materials (**Appendix 5**). Systolic pressure at three minutes was 127.54 ± 23.70 mmHg, change: -3.18 ± 16.80 mmHg, and at six minutes was 123.06 ± 22.28, change: -7.65 ± 17.27. Diastolic pressures at three minutes were 64.65 ± 13.56 mmHg, change: -4.65 ± 9.82 mmHg, and at 6 minutes 62.81 ± 12.53 mmHg, change: -6.49, 9.64 mmHg.

### Topographical maps of adenosine-evoked sensations

Bodily sensations were almost nonexistent in the baseline state, yet adenosine induced significant and topographically distinct sensation patterns (**Figure 1)**. When we directly compared sensations during adenosine with baseline state (**Figure 1**), we observed significant increases in sensations of pressure and pain around the chest area, sensations of warmth in the upper body, including the face. Sensations of weakness were less common, but they were also concentrated in the chest and torso areas. The cumulative sensation map revealed that adenosine-elicited sensations were most consistently felt in the torso, especially in the chest, shoulder, and head area. These effects were consistent for participants with and without myocardial ischemia, and direct comparisons between the ischemic and non-ischemic patients revealed only one significant cluster of pixels in the knee of the right leg in the bodily map of warmth.

**Figure 1.**
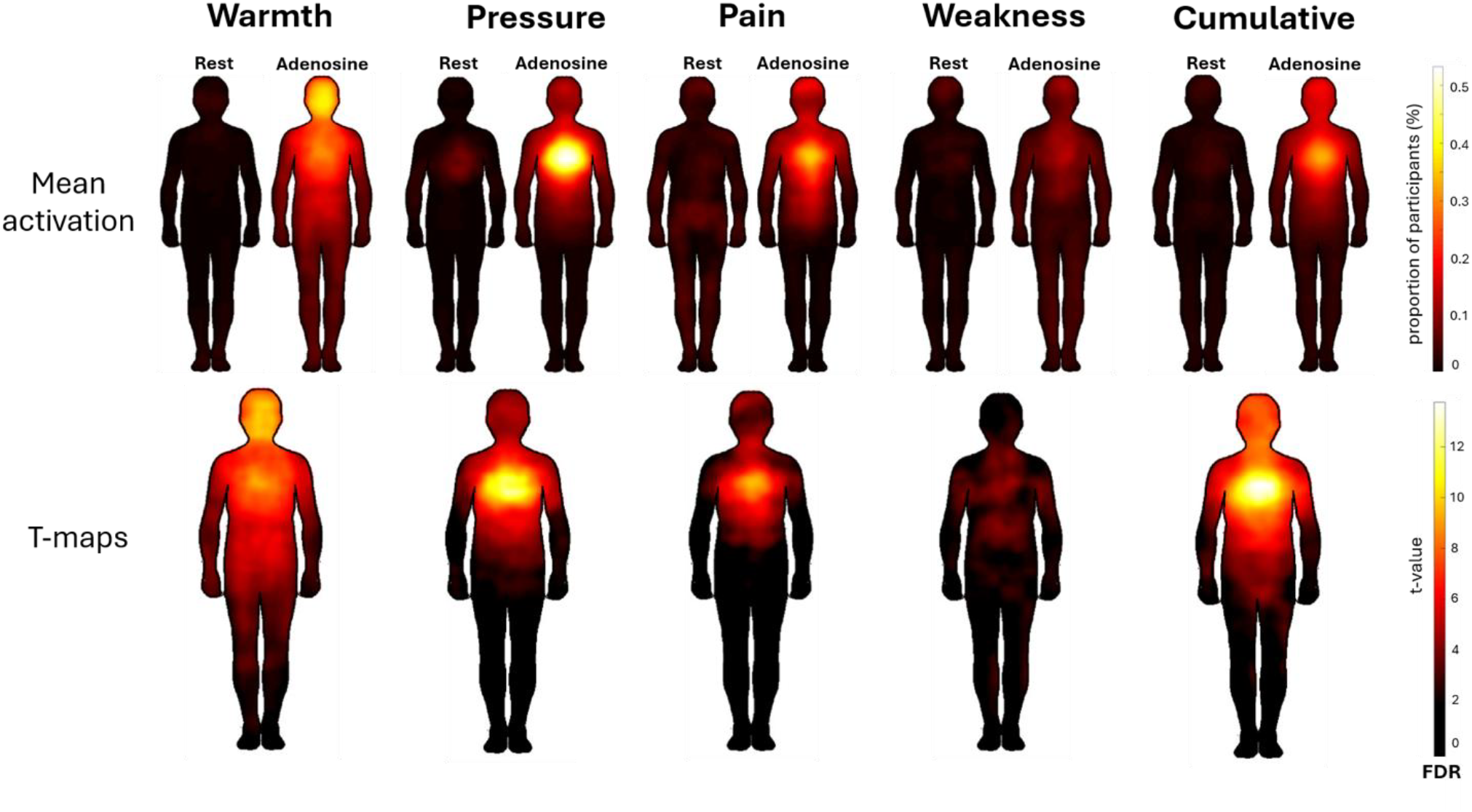
Topographical maps of bodily sensations during rest and stress (top) and the regions where sensations were stronger during adenosine versus rest (bottom). Data are thresholded at p < 0.05, FDR corrected.

### Emotions and subjective sensations elicited by adenosine

Adenosine evoked prominent emotional and somatic sensations in the participants (**Figure 2** and **Table S-1)**. Across the sample, adenosine induced an increase in all the measured somatic sensations, such as *rapid breathing, pressure in the chest, dizziness*, except for *fatigue* (*p*_FDR_ < 0.001), and in all negative emotions (e.g., anxiety and fear), as well as a decrease in positive emotions, namely *calmness, joy*, and *happiness* (*p*_FDR_ < 0.001). Changes in *anger* and *sadness* were not significant (*p*_FDR_ > 0.05).

**Figure 2.**
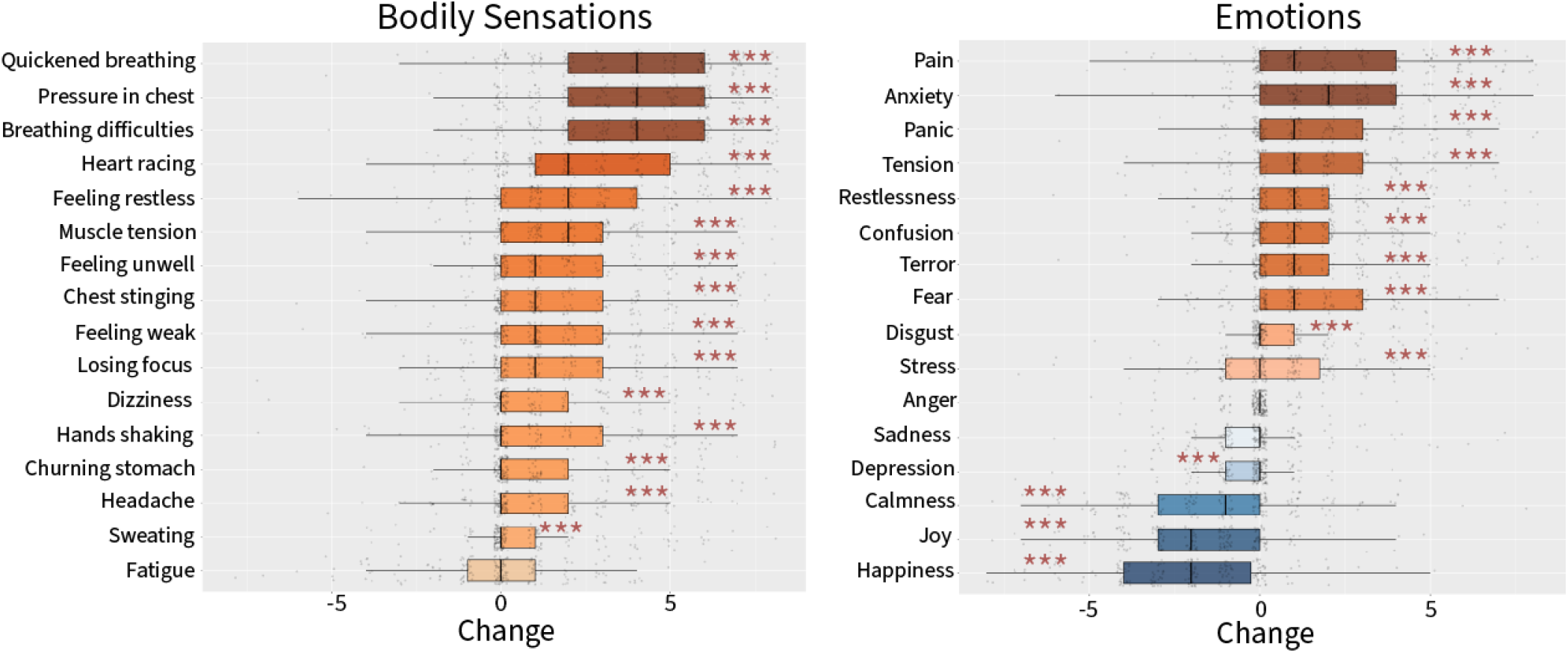
Adenosine induces A. bodily sensations and B. emotions. Each figure shows the distribution of change scores calculated across all participants, including both ischemic and non-ischemic subjects. A box plot represents the interquartile range of

### Emotions and sensations elicited by adenosine in patients with and without myocardial ischemia

Overall, the effects were consistent across sexes and ischemic versus non-ischemic patients, but we also observed some differences. The two-way aligned rank-transformed ANOVA revealed a statistically significant main effect of **ischemia** on *sweating* (*F*(1, 178) = 13.38, *p*_FDR_ = 0.023, 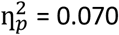), and the main effect of **sex** on sensations of *shaking hands* (*F*(1, 178) = 12.68, *p*_FDR_ = 0.023,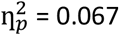) and *anger* (*F*(1, 178) = 9.94, *p*_FDR_ = 0.048,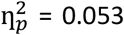). The ANOVA also showed a significant **interaction of ischemia and sex** on reported feeling of *panic* (*F*(1, 178) = 9.83, *p*_FDR_ = 0.048,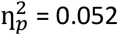).

We then performed a follow-up Wilcoxon aligned rank t-tests to assess the direction of differences in reported sensations and emotional responses between ischemic and non-ischemic, and between male and female subjects. Wilcoxon aligned rank t-test **across groups** revealed that non-ischemic patients reported stronger sensations of *muscle tension* (change mean ± SD; non-ischemic: 2.48 ± 2.03, ischemic: 1.55 ± 2.11, *W* = 5152.5, *p*_FDR_ = 0.041), and *panic* (non-ischemic: 2.20 ± 2.22, ischemic: 1.07 ± 2.00, *W* = 5213.0, *p*_FDR_ = 0.040 compared to ischemic patients. Wilcoxon t-test **across sexes** revealed that women reported stronger sensations of *shaking hands* (females: 1.91 ± 2.36, males: 0.63 ± 2.14, *W* = 5595.5, *p*_FDR_ = 0.00064), *heart racing* (females: 3.27 ± 2.35, males: 1.89 ± 2.51, *W* = 5498.0, *p*_FDR_ = 0.0019) and *muscle tension* (females: 2.50 ± 2.02, males: 1.61 ± 2.12, *W* = 5188.5, *p*_FDR_ = 0.030) compared to men.

### Associations between bodily sensation maps and the strength of emotional and bodily responses

We next investigated the relationship between the emotional and somatic responses to adenosine and the net sensations experienced in the body, as indicated by the bodily sensation maps. **Figure 3** illustrates the correlations between the net bodily sensations (i.e., total number of pixels colored in the body maps) and self-reported responses in 16 bodily sensations and 16 emotions (presented in Figure 2). Net sensation changes in the bodily maps significantly correlated with most of the self-reported bodily responses. *Pain, terror, panic, anxiety, restlessness*, and *tension* correlated with net changes in sensation across most, if not all, bodily maps. Other emotions like depression, sadness, anger, calmness, joy, and happiness correlated very weakly with sensations in all bodily maps.

**Figure 3.**
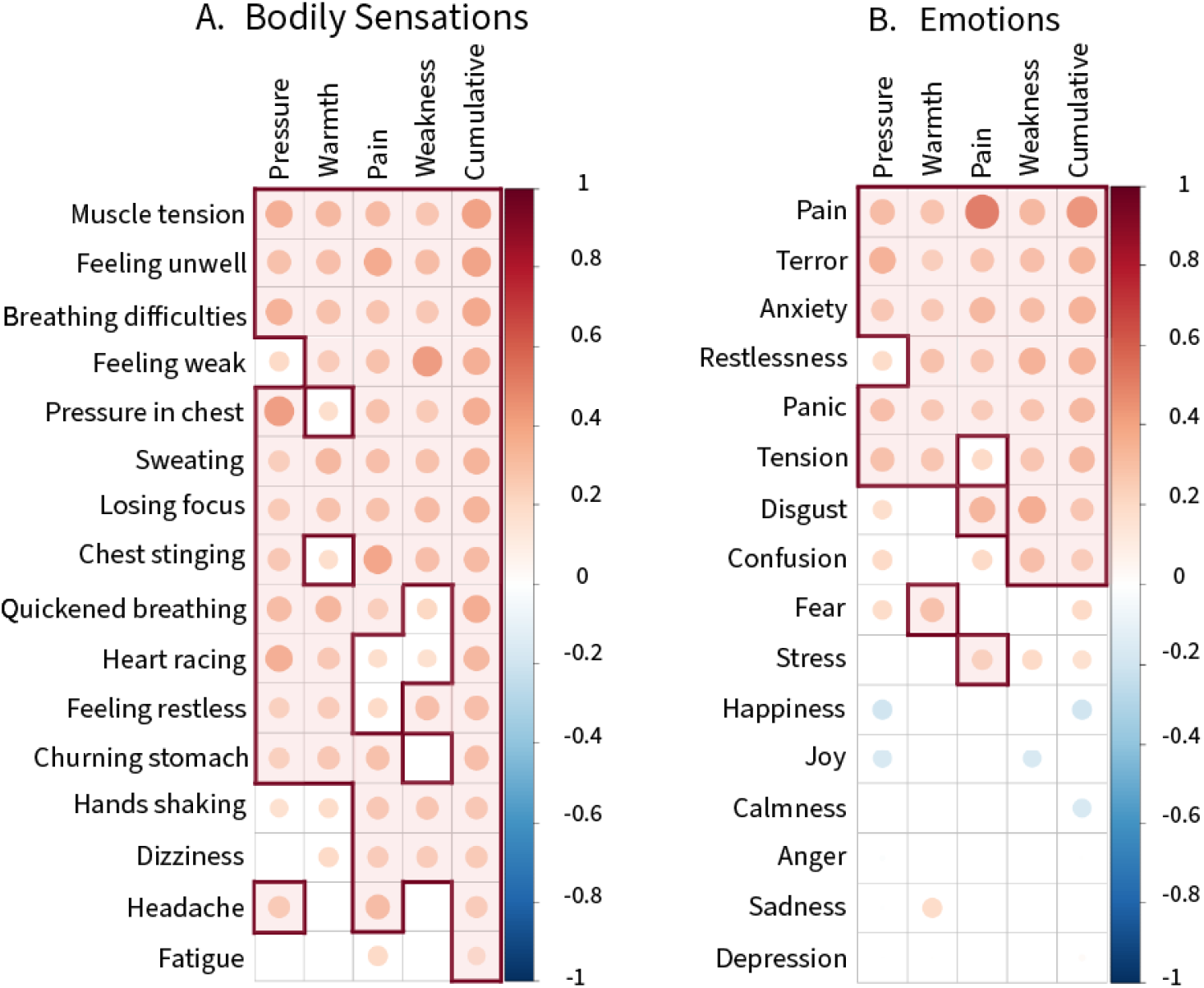
Correlations (pFDR < 0.05) net felt sensations changes in the body with (A.) bodily sensations and (B.) emotional responses elicited by adenosine. Correlations with Spearman’s rho > 0.20 are highlighted.

### Associations between hemodynamic responses and emotional and somatic sensations

Somatic sensations and emotions were also significantly associated with heart rate response. Higher peak HR response (from the baseline) was significantly associated with changes in several emotions, namely anxiety, *terror, fear, confusion, panic, sadness, anger*, and somatic sensations, such as *quickened breathing, breathing difficulties, muscle tension, heart racing, hands shaking, feeling restless*, and *sweating*, Regression coefficients presented in Figure 4 (detailed statistical results **in supplementary table S7**) can be interpreted as follows: each 10-bpm unit increase in highest HR elevation was associated with a change in subjective ratings equal to the respective beta estimate. For instance, in the case of sensation of quickened breathing, exhibiting peak HR response higher by 10 bpm is associated with a 0.62-point increase in the reported sensation (*b* = 0.62, 95% CI [0.29, 0.95], *p*_*FDR*_ = 0.0036, *β* = 0.34).

**Figure 4.**
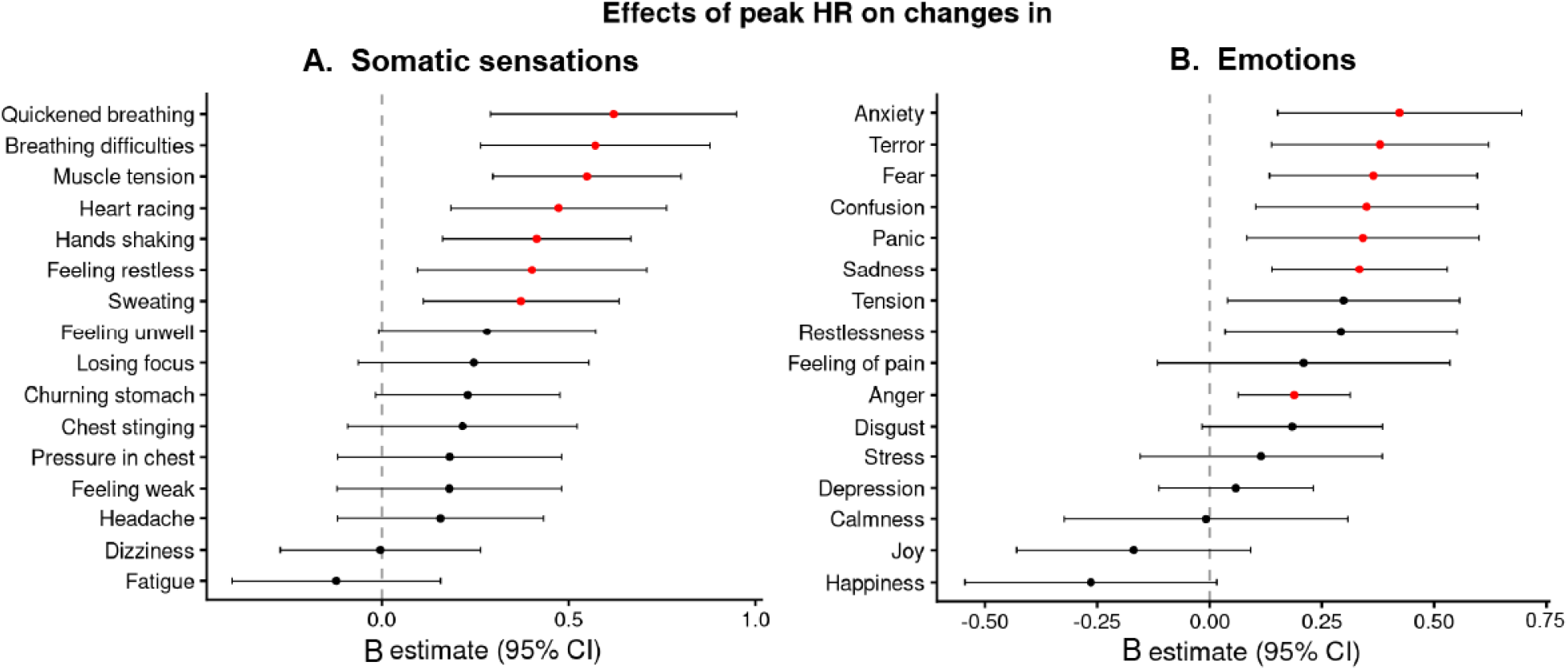
Effects of peak HR response on changes in rating of emotions and somatic sensations. Significant effects (p_FDR_ < 0.05) are highlighted in red.

Heart rate at the three-minute mark was a significant predictor of some of the same emotions and somatic responses, namely, *quickened breathing, muscle tension, breathing difficulties, heart racing, sweating, fear, confusion, shaking hands*, and *sadness*. However, blood pressure readings at three minutes were not good predictors, as diastolic pressure at three minutes explained only the variance in changes in *anxiety*. Detailed results can be found in Supplementary Materials tables S8, S9, and S10.

## Discussion

Our main finding was that peripheral physiological activation caused by adenosine leads to large-scale changes in bodily sensations, which translate into emotional experiences. Critically, these experiences were directly associated with systemic hemodynamic response to adenosine and occurred in the absence of direct pharmacological perturbation of the CNS, demonstrating the causal link between physiological changes and emotional experience. These responses are consistent between sexes as well as across patients with and without myocardial ischemia. All in all, our results suggest a generalizable link between physiological activation, autonomic signaling, interoception, and emotion (Nummenmaa et al., 2014).

### Somatic and emotional responses to adenosine

Adenosine induces sensations of dyspnea, chest pain, body tension, headache, and sensations attributable to peripheral vasodilation (Cerqueira et al., 1994), and carotid body chemoreceptor activation (Biaggioni et al., 1987), and increased sympathetic nerve traffic (Biaggioni et al., 1991). Using the topographical bodily mapping tool, we demonstrated the spatially distinct topography of the somatic sensations. Adenosine-induced experience of pain and pressure mostly in the chest area, while experiences of warmth were evoked throughout the body, weakening in the distal limbs. Feelings of weakness were less salient and mostly experienced in the torso. To our knowledge, this is the most detailed description of the subjective experiences caused by adenosine that go beyond previously well-established such as sensations of dyspnea, flushing, headache, gastrointestinal discomfort, chest pain, and light-headedness (Burki et al., 2005; Cerqueira et al., 1994).

These somatic sensations were associated with a prominent shift in the participants’ emotional state. In general, adenosine increased the experience of all negative emotions (most notably pain, anxiety, and panic) except for anger and sadness, while decreasing the experience of positive emotions such as joy and happiness. The emotional experiences (most notably pain, terror, anxiety, panic, disgust, tension, and restlessness) correlated with the magnitude of the topographical localization warmth, pain, pressure, and weakness, indicating a linkage between autonomic interoception and emotions.

### Association of psychological self-reports with hemodynamic measurements

Reflexive increase in heart rate due to systemic drop in vasodilation and chemoreceptor stimulation (Biaggioni et al., 1987, 1991; Timmers et al., 2004) consistently predicted the strength of somatic sensations and emotions. Peak heart rate response was associated with somatic experiences of **muscle tension, heart racing, breathing difficulties, quickened breathing, sweating, shaking hands**, and **feeling restless**, and emotional responses of **fear, terror, panic, anxiety, confusion, sadness, and anger**, when controlled for sex and ischemic status. This association aligns with the view that physiological cues can shift the emotional states (Barrett & Lindquist, 2008; Critchley & Harrison, 2013). We demonstrate that this occurs even when the peripheral physiological activation caused by vasodilatation is triggered without a direct pharmacological action in the CNS.

The observed lack of association between somatic sensations mentioned above and blood pressure changes can be explained by the distinct neural and secondary origins of these sensations. Subjective sensations related to shortness of breath are predominantly mediated by activation of adenosine receptors in the carotid bodies (Biaggioni et al., 1987). Stimulation of these chemoreceptors has been shown to drive the increase in heart rate during adenosine administration (Biaggioni et al., 1987; Timmers et al., 2004) and it explains the observed association between sensation of shortness of breath and changes in heart rate, independent of changes in blood pressure. Sensations of **muscle tension** and **shaking hands** could be attributed to increased muscle sympathetic nerve traffic observed during adenosine administration (Biaggioni et al., 1991; Engelstein et al., 1994). The increase in the sympathetic traffic is not mediated by chemoreceptor activation, but is an afferent consequence of systemic cardiovascular effects of adenosine, parallel to the increase in heart rates (Engelstein et al., 1994).

### Myocardial ischemia is not associated with the subjective sensations of adenosine

Patients with myocardial ischemia exhibit a blunted cardiac response to pharmacological vasodilator stressors (Gorur et al., 2012; Tomiyama et al., 2015), which might reflect impairments in baroreceptor activity and the autonomic system (Abidov et al., 2003). We show that the heart rate reflex was weaker in ischemic than in non-ischemic patients. However, having stress-induced ischemia in the myocardium did not alter sensitivity to the topographical organization or intensity of somatic sensations or the concomitantly arising emotions. Instead, we found no strong evidence for ischemic myocardial pathology influencing adenosine-induced emotions and sensations. Despite marked peak HR differences across the groups, this difference did not translate to markedly different subjective sensations between ischemic and non-ischemic patients. Additional factors beyond myocardial ischemia must be investigated to explain why the physiological differences in HR response did not translate into different subjective experiences across the clinical groups.

Finally, the subjective sensations and emotions were generally consistent across sexes. This occurred despite female patients showing higher heart rates than males, and we found sex differences only for a limited number of somatic sensations, such as **shaking hands, heart racing**, and **muscle tension**. We did not find any sex differences in emotional experiences. These data underline the well-established sex-invariant emotional processing, which is manifested here despite the markedly differential physiological responses across sexes (Volynets et al., 2020)

### Limitations

All our participants were patients with suspected myocardial ischemia, although the myocardial perfusion imaging only confirmed this in a subset of the studied individuals. Since all participants were symptomatic patients referred to evaluation of the presence of functionally significant coronary artery disease, the heart-brain connection and cardioceptive integration may already be strongly shifted across our sample. However, our results show that symptomatic myocardial ischemia does not influence sensitivity to somatically induced emotional responses.

Future studies should nevertheless replicate the effect in healthy control subjects. Myocardial ischemia was also unequally distributed across sexes. However, sex differences in all the dependent variables were mostly non-existent, and we controlled for the imbalance statistically in the main analyses. Finally, we lack detailed data on adherence to the long-term use of the medication. The effect of cardiovascular medication on affective disorders would be important to understand in the context of clinical studies.

## Conclusions

Adenosine-induced peripheral physiological and subsequent autonomic activation evokes a substantial increase in somatic sensations, which in turn are predictive of the most prominent negative emotional changes. These results emphasize how subjective emotional states are brought about by the interoception of autonomic activity. Because we utilized only a single pharmacological stimulus, leading to large-scale autonomic activity, leading to a predominantly negative emotional state, future studies need to establish how distinct patterns of autonomic activation are associated with differentiated emotional experiences.

## Abbreviations

CVD: cardiovascular disease
CT: computed tomography
FDR: false discovery rate
pCST: pharmacological cardiac stress test
PET: positron emission tomography

## Acknowledgements

This work was supported by European Research Council (advanced grant #101141656 to LN). The funders had no role in designing or reporting the study.

