## Supplementary Materials for "Emotions and bodily sensations evoked by cardiac adenosine stress"

### Appendix 1. Bodymaps of adenosine-induced sensations

#### 1.1. Descriptive statistics on the bodily maps

Table S1 shows the number of pixels colored in each topographical bodily map in both rest and adenosine stress, as well as the change in the number of pixels (stress - rest).

*Table S1. Number of pixels colored in each bodily map (warmth, pressure, pain, and weakness) in both rest and after adenosine stress. The values are expressed in the form mean ± SD.*

| Bodymap | Rest | Adenosine | Change |
|---|---|---|---|
| *Cumulative* | 1139.94 ± 2243.58 | 5290.07 ± 6323.54 | 4150.13 ± 5988.37 |
| *Weakness* | 908.26 ± 2826.99 | 3310.13 ± 8675.50 | 2401.87 ± 8796.48 |
| *Pain* | 2109.48 ± 4887.00 | 4818.12 ± 6909.46 | 2708.64 ± 6476.84 |
| *Pressure* | 827.35 ± 1638.68 | 5999.09 ± 8074.70 | 5171.74 ± 7728.99 |
| *Warmth* | 625.75 ± 3116.89 | 8080.11 ± 11671.91 | 7454.36 ± 11665.99 |

#### 1.2. The adenosine response maps

We created bodily maps illustrating the change of sensations induced by adenosine from rest by subtracting the activations reported in rest from the adenosine stress bodily maps. These mean response bodily maps are illustrated in **Figure S1**. Since these change bodily maps are average difference maps across subjects, the pixel intensity illustrates the proportion of subjects that reported a change in the specific pixel.

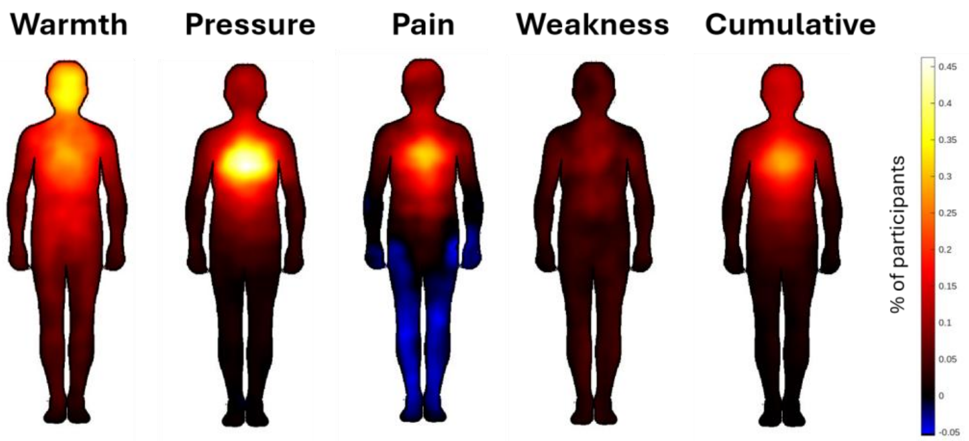


*Figure S1. Baseline normalized mean localization of four sensations in the body (warmth, pressure, pain, and weakness) induced by adenosine.*

#### 1.3. Group difference topographical maps

Table S2 shows the differences in the number of colored pixels across ischemic and non-ischemic patients at rest and during adenosine stress, as well as the subtracted change.

*Table S2. Number of pixels colored in each bodily map (warmth, pressure, pain, and weakness) in both rest and after adenosine stress across ischemic and non-ischemic patients. The values are expressed in form mean ± SD.*

| | Rest | | Adenosine | | Change | |
|---|---|---|---|---|---|---|
| **Ischemia** | **No** | **Yes** | **No** | **Yes** | **No** | **Yes** |
| *Cumulative* | 984.03 ± 1668.35 | 1327.83 ± 2784.14 | 5584.81 ± 6577.31 | 4934.86 ± 6026.49 | 4600.78 ± 6520.44 | 3607.03 ± 5266.99 |
| *Weakness* | 900.89 ± 3023.90 | 917.06 ± 2590.51 | 3748.44 ± 9689.54 | 2786.29 ± 7306.29 | 2847.55 ± 10147.26 | 1869.23 ± 6867.49 |
| *Pain* | 1857.08 ± 3521.17 | 2405.49 ± 6124.61 | 4624.18 ± 6652.64 | 5045.58 ± 7234.15 | 2767.09 ± 6982.99 | 2640.09 ± 5870.28 |
| *Pressure* | 793.26 ± 1474.91 | 869.01 ± 1827.63 | 6894.90 ± 9303.91 | 4904.22 ± 6135.21 | 6101.64 ± 8901.31 | 4035.21 ± 5850.07 |
| *Flushing* | 306.54 ± 897.41 | 1012.80 ± 4515.31 | 9347.73 ± 13645.22 | 6543.11 ± 8537.20 | 9041.20 ± 13608.56 | 5530.31 ± 8444.79 |

#### 1.4. Mean topographical bodily maps for patients with stress-induced myocardial ischemia versus those without

In Figure S2, we illustrated the mean adenosine-induced topographical activation separately for patients with cardiac stress-induced myocardial ischemia and without.

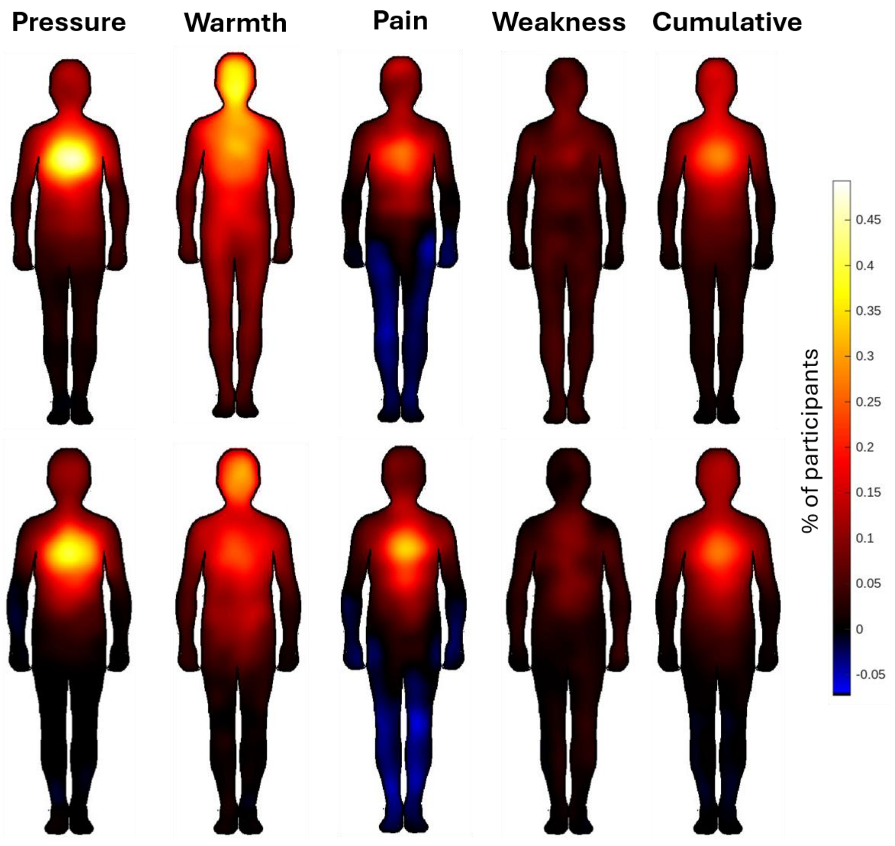


*Figure S2. The figure shows the consistency of localization of four sensations in the body (warmth, pressure, pain, and weakness) at rest and during adenosine stress for non-ischemic (top) and ischemic (bottom) patients. The figure includes a map of all these sensations averaged out into a cumulative map (top figure).*

#### 1.5. Mean difference in bodily maps between ischemic and non-ischemic patients

We illustrated the differences in mean adenosine responses between patients with stress-induced myocardial ischemia and those without in Figure S3.

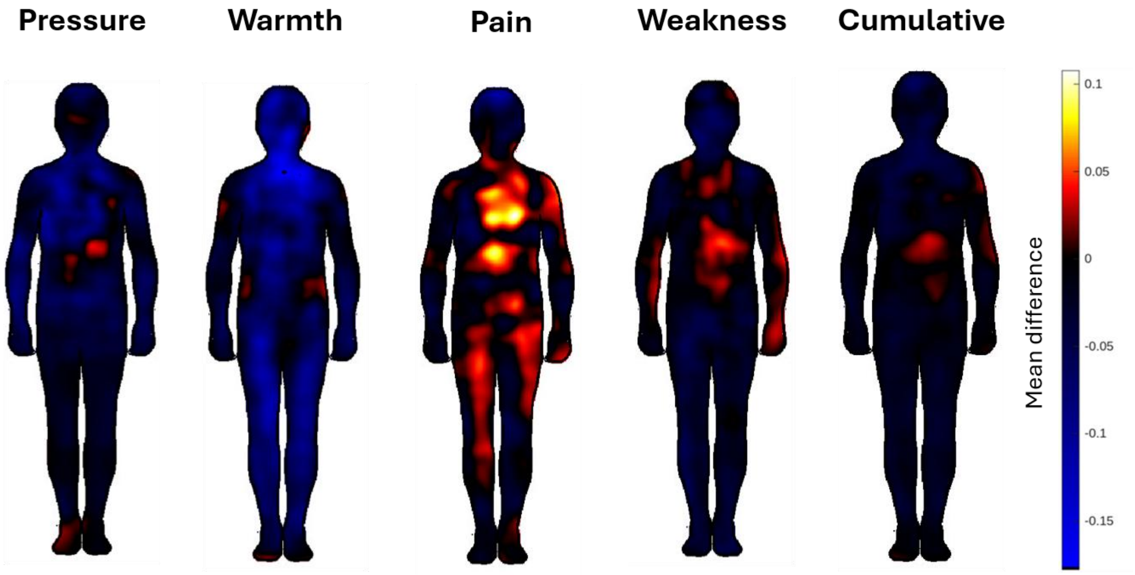


*Figure S3. Bodily maps of differences in the topographical localization of sensations between ischemic and non-ischemic subjects. Topographical difference maps were computed by pixel-wise subtracting the non-ischemic mean response from the ischemic mean response.*

#### 1.6 Parametric difference t-maps between ischemic and non-ischemic patients

We conducted a pixelwise univariate t-test to investigate statistically significant differences between ischemic and non-ischemic patients. Statistically significant (after FDR-correction) pixels are highlighted in Figure S4.

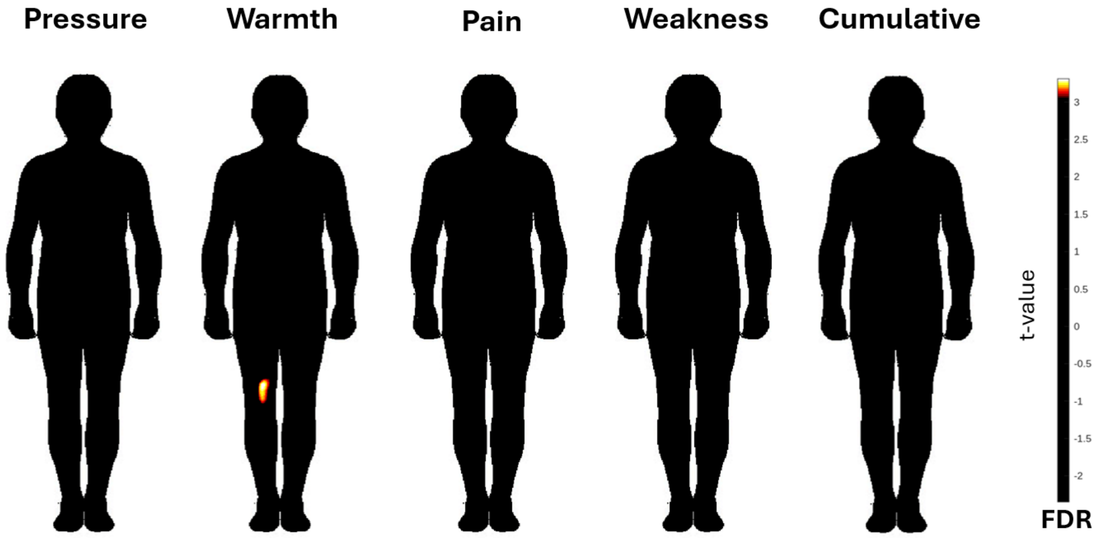


*Figure S4. Parametric t-maps of the statistically significant differences between ischemic and non-ischemic patients. All significant t-values are colored non-black.*

### Appendix 2. Associative bodily maps

To investigate the associations between topographical localization of sensations and the strength of bodily and emotional responses induced by adenosine, we performed pixel-wise linear regression analyses. We computed separate, pixel-wise, simple linear regression models for each bodily map (**Figures S5-8**) modality (cumulative, pressure, weakness, pain, warmth) using change in experience bodily sensation and emotion scores as predictors. We estimated regression coefficients, t-values, and two-tailed p-values using matrix-based ordinary least squares regression implemented with QR decomposition in MATLAB. We thresholded the beta-weights using the FDR-corrected p-value as the threshold. We had bodily maps from 181 participants.

#### 2.1 Cumulative association maps

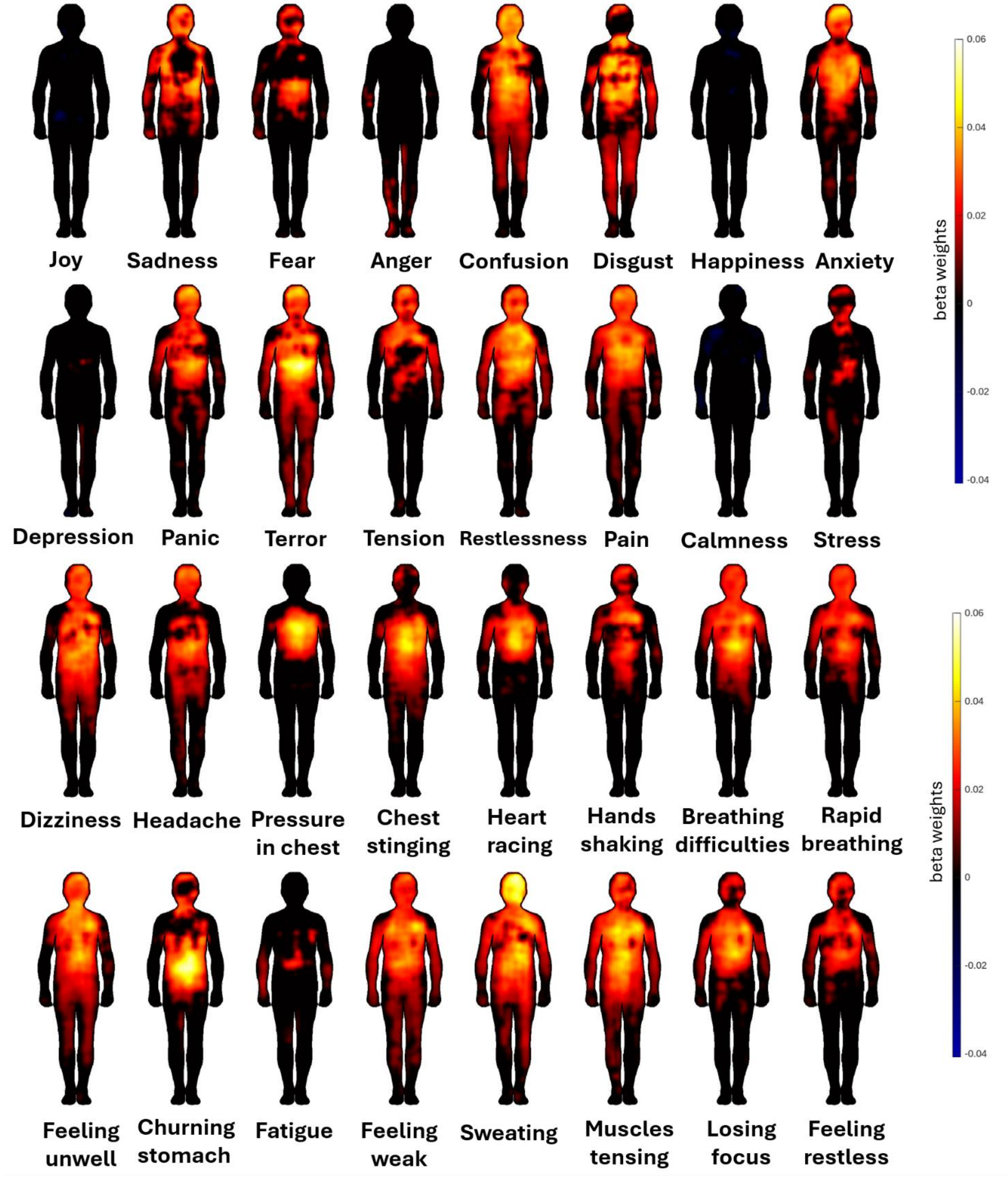


*Figure S5. Associations of change in 16 emotions and 16 bodily sensations with the cumulative sensations bodily map.*

#### 2.2. Pressure association maps

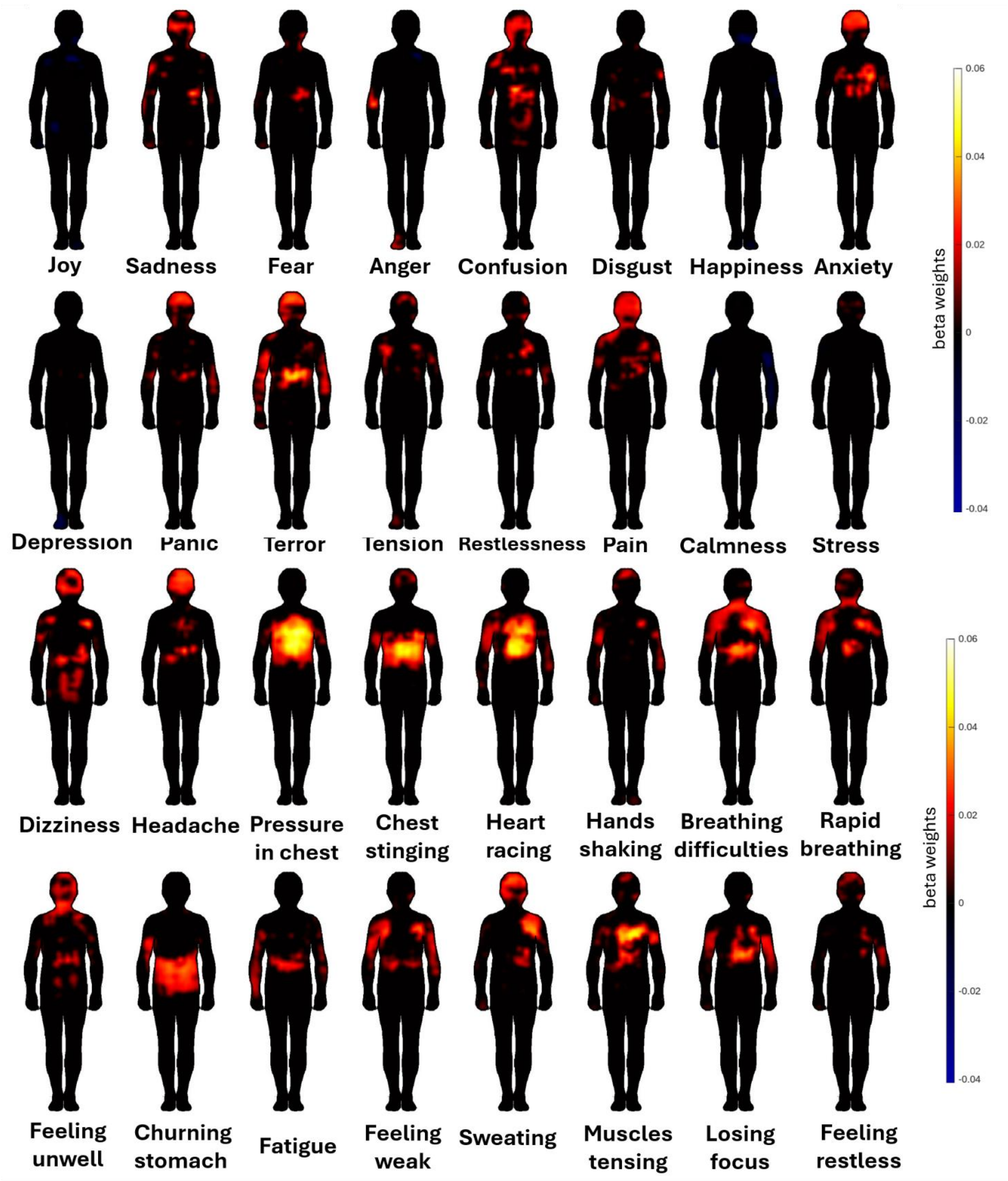


*Figure S6. Associations of change in 16 emotions and 16 bodily sensations with topographical maps of sensation of pressure.*

#### 2.3. Weakness association maps

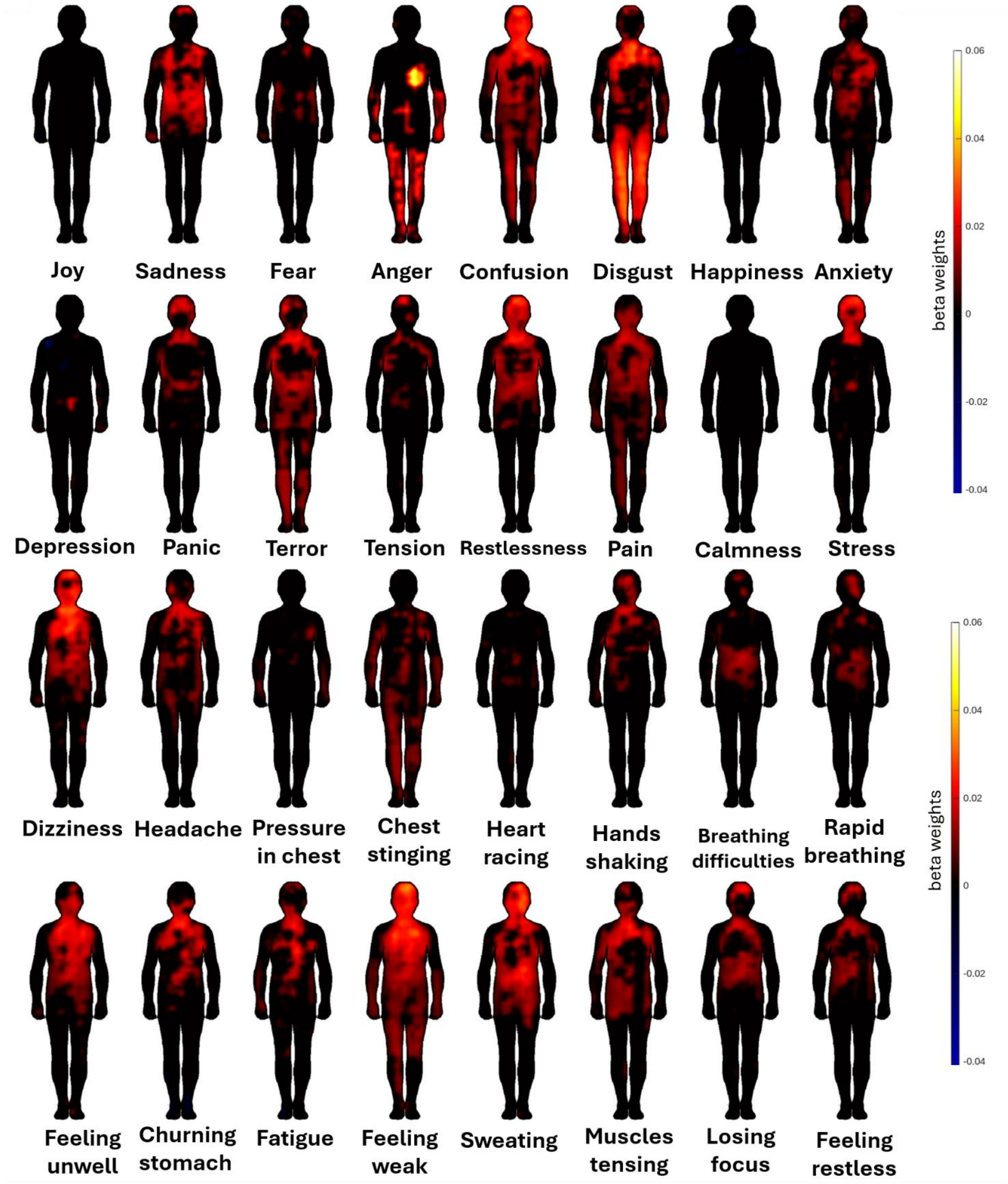


*Figure S7. Associations of change in 16 emotions and 16 bodily sensations with topographical maps of sensation of weakness.*

#### 2.4. Pain association maps

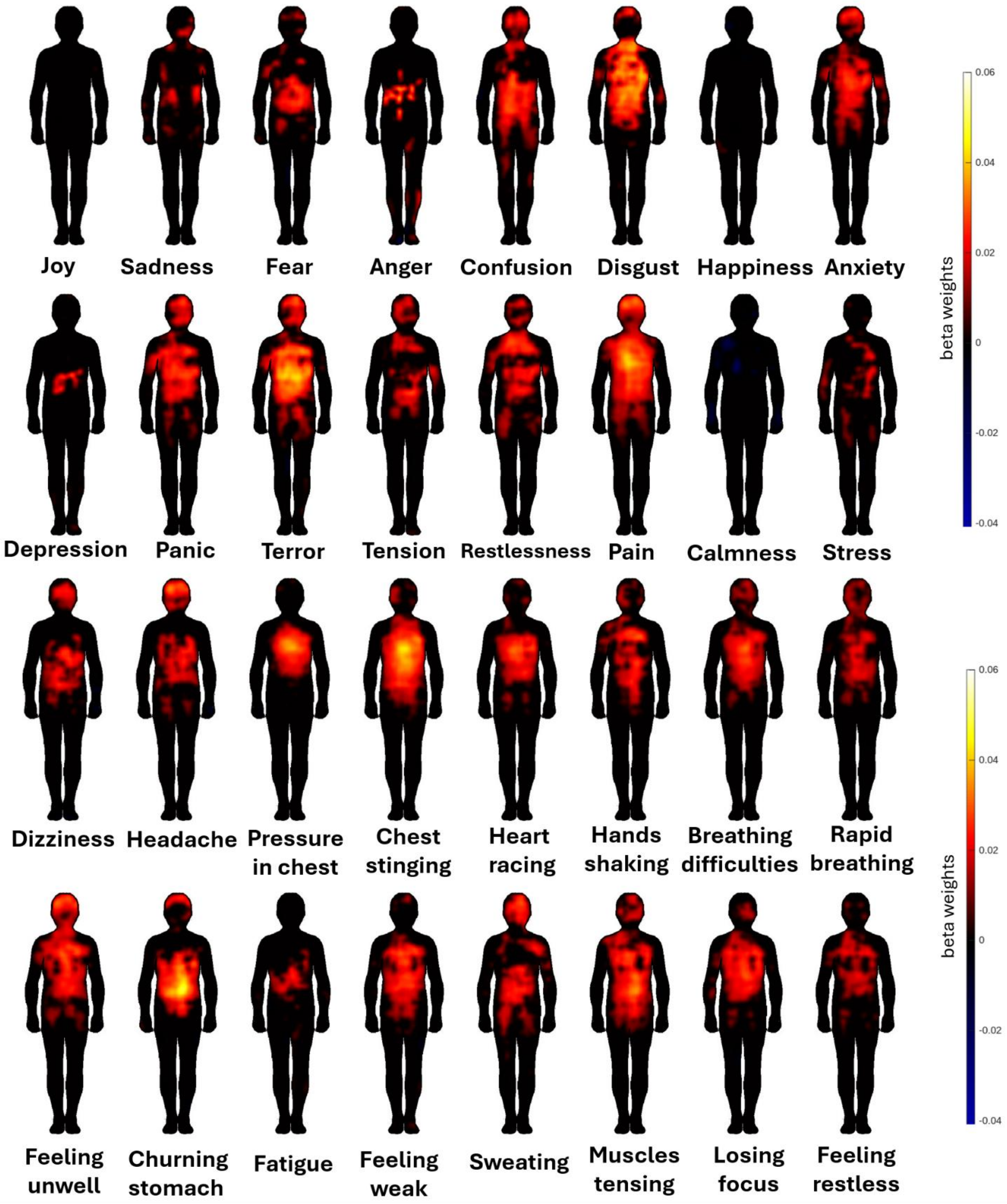


*Figure S8. Associations of change in 16 emotions and 16 bodily sensations with topographical maps of sensation of pain.*

#### 2.5 Warmth association maps

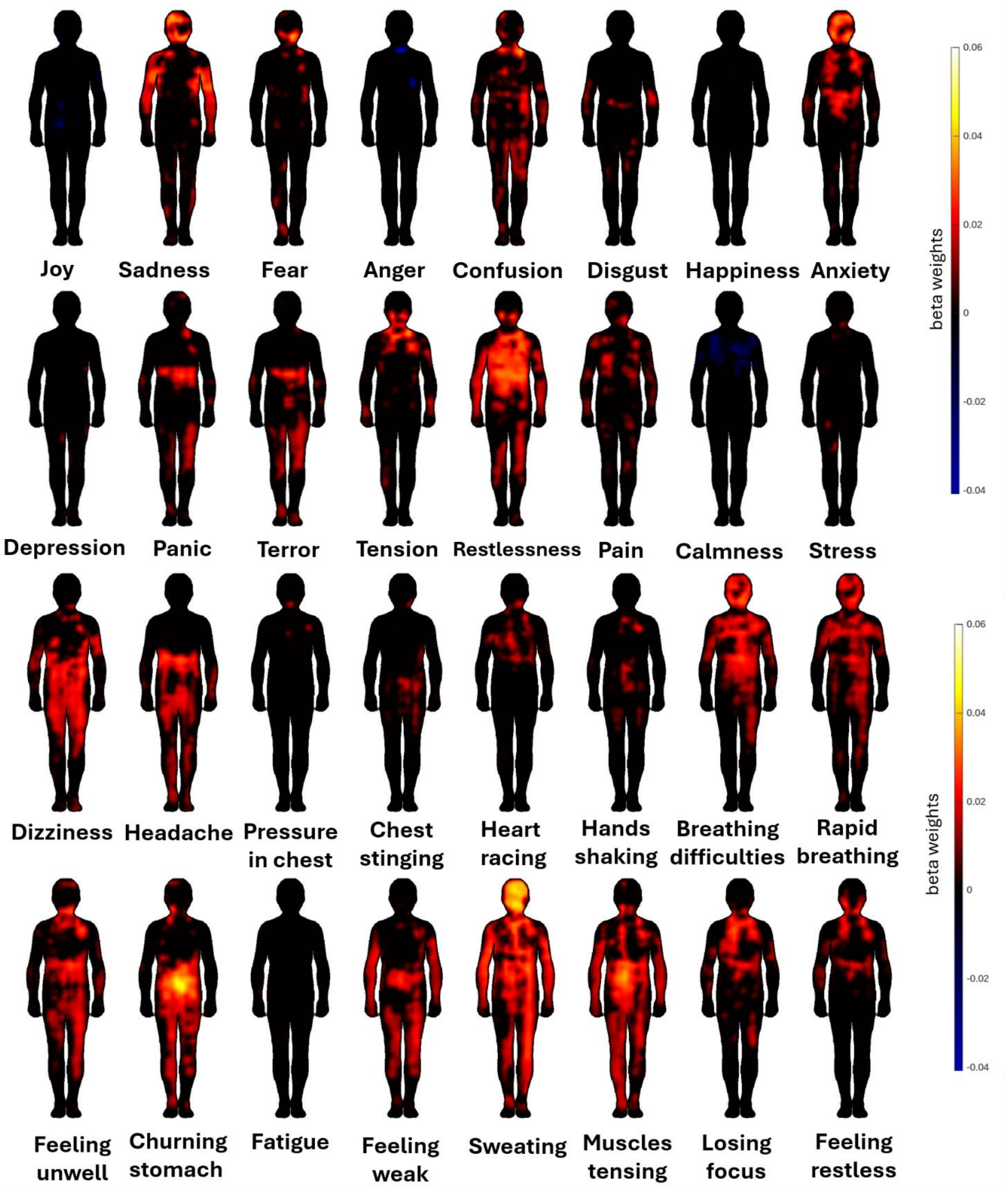


*Figure S9. Associations of change in 16 emotions and 16 bodily sensations with topographical maps of sensation of warmth.*

### Appendix 3. Descriptive statistics and detailed statistical results on self-reports of 16 emotions and 16 bodily sensations

#### 3.1 Descriptive statistics and statistical results across all subjects

Here we collected descriptive statistics of self-reports of 16 emotions and 16 bodily sensations. **Table S3** contains mean scores from rest, adenosine stress, and change scores across all subjects, as well as p-values and W-value of the Wilcoxon t-test between rest and adenosine stress conditions.

*Table S3. Table of descriptive statistics (mean ± SD) and statistical results (W- and p-values) of emotions and bodily sensation scores at rest and during adenosine stress, as well as change scores.*

| Psychological response | **Rest** | **Adenosine** | **Change** | **p-value** | **pFDR** | **W-value** |
|---|---|---|---|---|---|---|
| *Joy* | 4.43 ± 1.98 | 2.63 ± 2.09 | -1.81 ± 2.06 | 3.39E-19 | 1.36E-18 | 9919.5 |
| *Sadness* | 2.08 ± 1.44 | 1.97 ± 1.69 | -0.12 ± 1.72 | 0.083 | 0.088 | 2792.5 |
| *Fear* | 2.61 ± 1.79 | 3.84 ± 2.37 | 1.23 ± 1.96 | 6.49E-13 | 1.04E-12 | 1134 |
| *Anger* | 1.28 ± 0.91 | 1.21 ± 0.67 | -0.08 ± 0.91 | 0.36 | 0.38 | 453 |
| *Confusion* | 1.81 ± 1.16 | 3.17 ± 2.17 | 1.35 ± 2.06 | 2.89E-15 | 5.77E-15 | 624 |
| *Disgust* | 1.23 ± 0.79 | 1.96 ± 1.65 | 0.72 ± 1.66 | 1.08E-08 | 1.32E-08 | 247 |
| *Happiness* | 5.00 ± 2.12 | 2.92 ± 2.23 | -2.10 ± 2.21 | 5.40E-21 | 2.47E-20 | 10522.5 |
| *Anxiety* | 2.26 ± 1.61 | 4.40 ± 2.38 | 2.15 ± 2.27 | 1.73E-21 | 9.25E-21 | 514.5 |
| *Depression* | 2.01 ± 1.49 | 1.58 ± 1.27 | -0.44 ± 1.29 | 2.77E-06 | 3.17E-06 | 2569.5 |
| *Panic* | 1.46 ± 1.05 | 3.15 ± 2.33 | 1.69 ± 2.19 | 3.12E-18 | 9.97E-18 | 307.5 |
| *Terror* | 1.33 ± 0.91 | 2.63 ± 2.19 | 1.31 ± 2.06 | 8.08E-15 | 1.52E-14 | 303 |
| *Tension* | 2.86 ± 1.68 | 4.45 ± 2.37 | 1.57 ± 2.08 | 4.36E-17 | 1.07E-16 | 1131.5 |
| *Restlessness* | 2.32 ± 1.59 | 3.69 ± 2.32 | 1.37 ± 2.15 | 1.95E-14 | 3.47E-14 | 910 |
| *Pain* | 1.83 ± 1.45 | 3.96 ± 2.60 | 2.16 ± 2.67 | 4.96E-18 | 1.32E-17 | 570.5 |
| *Calmness* | 4.57 ± 2.36 | 3.27 ± 2.05 | -1.24 ± 2.57 | 1.98E-09 | 2.54E-09 | 7903 |
| *Stress* | 3.00 ± 1.84 | 3.58 ± 2.24 | 0.57 ± 2.06 | 0.00011 | 0.00012 | 2670.5 |
| *Dizziness* | 1.48 ± 0.96 | 2.78 ± 2.15 | 1.29 ± 2.16 | 2.71E-13 | 4.57E-13 | 495 |
| *Headache* | 2.18 ± 1.57 | 3.29 ± 2.41 | 1.13 ± 2.23 | 8.08E-10 | 1.08E-09 | 846 |
| *Pressure in chest* | 1.59 ± 1.13 | 5.32 ± 2.49 | 3.75 ± 2.48 | 2.52E-27 | 4.03E-26 | 37 |
| *Chest stinging* | 1.38 ± 1.06 | 3.08 ± 2.31 | 1.70 ± 2.32 | 1.15E-16 | 2.64E-16 | 404.5 |
| *Heart racing* | 2.08 ± 1.48 | 4.65 ± 2.67 | 2.59 ± 2.52 | 6.15E-23 | 4.92E-22 | 450.5 |
| *Hands shaking* | 1.69 ± 1.40 | 2.97 ± 2.37 | 1.28 ± 2.34 | 1.46E-11 | 2.03E-11 | 682 |
| *Breathing difficulties* | 1.65 ± 1.13 | 5.26 ± 2.51 | 3.63 ± 2.58 | 1.48E-27 | 4.03E-26 | 200 |
| *Quickened breathing* | 1.71 ± 1.06 | 5.61 ± 2.51 | 3.91 ± 2.67 | 4.58E-27 | 4.89E-26 | 218 |
| *Feeling unwell* | 1.22 ± 0.55 | 3.10 ± 2.53 | 1.90 ± 2.41 | 3.52E-18 | 1.02E-17 | 89.5 |
| *Churning stomach* | 1.28 ± 0.72 | 2.49 ± 2.19 | 1.21 ± 2.10 | 2.14E-12 | 3.11E-12 | 335 |
| *Fatigue* | 2.71 ± 1.82 | 2.58 ± 1.88 | -0.14 ± 2.18 | 0.59 | 0.59 | 3835 |
| *Feeling weak* | 1.68 ± 1.20 | 3.34 ± 2.35 | 1.65 ± 2.41 | 3.33E-16 | 7.10E-16 | 724 |
| *Sweating* | 1.45 ± 0.97 | 2.36 ± 2.08 | 0.90 ± 2.08 | 3.31E-08 | 3.93E-08 | 539 |
| *Muscle tension* | 2.11 ± 1.30 | 4.17 ± 2.37 | 2.06 ± 2.11 | 3.93E-22 | 2.52E-21 | 396.5 |
| *Losing focus* | 1.87 ± 1.37 | 3.32 ± 2.41 | 1.44 ± 2.40 | 1.18E-12 | 1.80E-12 | 933 |
| *Feeling restless* | 2.16 ± 1.54 | 4.24 ± 2.53 | 2.07 ± 2.48 | 8.12E-19 | 2.89E-18 | 826 |

### Appendix 4. Self-reported bodily and emotional responses across patients with stress-induced myocardial ischemia and without

We collected self-reports of 16 different bodily sensations and 16 emotions at rest and during adenosine stress. Here are the descriptive statistics of the responses stratified by the ischemia status.

#### 4.1. Descriptive statistics on bodily sensations and emotions in ischemic and non-ischemic subjects

Descriptive statistics on collected bodily sensations and emotions are presented separately for patients with cardiac stress-induced ischemia and without in **Table S4**.

*Table S4. Descriptive statistics (mean ± SD) on self-reported emotions and bodily sensations collected using a Likert scale at rest and during adenosine stress. The table also shows change scores (stress – rest).*

| | Rest | | Adenosine | | Change | |
|---|---|---|---|---|---|---|
| **Myocardial ischemia** | **No** | **Yes** | **No** | **Yes** | **No** | **Yes** |
| *Joy* | 4.63 ± 2.04 | 4.18 ± 1.88 | 2.53 ± 1.98 | 2.74 ± 2.23 | -2.14 ± 1.97 | -1.40 ± 2.11 |
| *Sadness* | 2.02 ± 1.45 | 2.15 ± 1.44 | 1.95 ± 1.68 | 2.00 ± 1.71 | -0.09 ± 1.62 | -0.15 ± 1.84 |
| *Fear* | 2.73 ± 1.77 | 2.45 ± 1.81 | 4.31 ± 2.37 | 3.27 ± 2.24 | 1.55 ± 2.12 | 0.83 ± 1.68 |
| *Anger* | 1.22 ± 0.71 | 1.36 ± 1.12 | 1.12 ± 0.38 | 1.33 ± 0.89 | -0.11 ± 0.78 | -0.04 ± 1.06 |
| *Confusion* | 1.71 ± 0.95 | 1.93 ± 1.37 | 3.24 ± 2.27 | 3.09 ± 2.05 | 1.52 ± 2.10 | 1.15 ± 2.00 |
| *Disgust* | 1.12 ± 0.48 | 1.37 ± 1.04 | 2.03 ± 1.78 | 1.87 ± 1.47 | 0.91 ± 1.82 | 0.49 ± 1.42 |
| *Happiness* | 5.12 ± 2.18 | 4.85 ± 2.06 | 2.91 ± 2.23 | 2.93 ± 2.25 | -2.26 ± 2.32 | -1.90 ± 2.07 |
| *Anxiety* | 2.18 ± 1.57 | 2.36 ± 1.67 | 4.65 ± 2.45 | 4.10 ± 2.27 | 2.49 ± 2.45 | 1.74 ± 1.96 |
| *Depression* | 1.80 ± 1.17 | 2.27 ± 1.79 | 1.42 ± 0.93 | 1.78 ± 1.57 | -0.37 ± 1.06 | -0.52 ± 1.53 |
| *Panic* | 1.40 ± 0.90 | 1.52 ± 1.21 | 3.58 ± 2.47 | 2.61 ± 2.02 | 2.20 ± 2.22 | 1.07 ± 2.00 |
| *Terror* | 1.34 ± 0.81 | 1.32 ± 1.02 | 2.94 ± 2.37 | 2.26 ± 1.91 | 1.62 ± 2.15 | 0.93 ± 1.88 |
| *Tension* | 2.99 ± 1.64 | 2.70 ± 1.73 | 4.83 ± 2.47 | 3.98 ± 2.18 | 1.82 ± 2.20 | 1.26 ± 1.89 |
| *Restlessness* | 2.28 ± 1.62 | 2.38 ± 1.57 | 3.98 ± 2.42 | 3.34 ± 2.16 | 1.70 ± 2.21 | 0.96 ± 2.02 |
| *Pain* | 1.82 ± 1.39 | 1.85 ± 1.52 | 4.36 ± 2.62 | 3.48 ± 2.51 | 2.56 ± 2.78 | 1.67 ± 2.45 |
| *Calmness* | 4.94 ± 2.21 | 4.12 ± 2.48 | 3.17 ± 2.04 | 3.40 ± 2.07 | -1.68 ± 2.42 | -0.71 ± 2.66 |
| *Stress* | 2.97 ± 1.76 | 3.04 ± 1.95 | 3.72 ± 2.32 | 3.40 ± 2.15 | 0.74 ± 1.99 | 0.35 ± 2.13 |
| *Dizziness* | 1.51 ± 1.08 | 1.44 ± 0.78 | 2.97 ± 2.29 | 2.54 ± 1.94 | 1.45 ± 2.32 | 1.09 ± 1.93 |
| *Headache* | 2.24 ± 1.66 | 2.10 ± 1.47 | 3.56 ± 2.52 | 2.95 ± 2.24 | 1.31 ± 2.22 | 0.90 ± 2.24 |
| *Pressure in chest* | 1.51 ± 1.21 | 1.68 ± 1.01 | 5.51 ± 2.47 | 5.09 ± 2.51 | 4.02 ± 2.50 | 3.41 ± 2.42 |
| *Chest stinging* | 1.48 ± 1.28 | 1.26 ± 0.68 | 3.14 ± 2.22 | 3.01 ± 2.43 | 1.66 ± 2.31 | 1.74 ± 2.35 |
| *Heart racing* | 2.27 ± 1.63 | 1.85 ± 1.25 | 5.25 ± 2.61 | 3.91 ± 2.58 | 3.01 ± 2.56 | 2.07 ± 2.38 |
| *Hands shaking* | 1.59 ± 1.19 | 1.81 ± 1.62 | 3.20 ± 2.34 | 2.70 ± 2.39 | 1.60 ± 2.32 | 0.89 ± 2.31 |
| *Breathing difficulties* | 1.63 ± 1.19 | 1.67 ± 1.06 | 5.58 ± 2.47 | 4.87 ± 2.52 | 3.98 ± 2.54 | 3.20 ± 2.57 |
| *Quickened breathing* | 1.74 ± 1.20 | 1.67 ± 0.87 | 5.98 ± 2.40 | 5.16 ± 2.59 | 4.26 ± 2.69 | 3.48 ± 2.59 |
| *Feeling unwell* | 1.15 ± 0.41 | 1.30 ± 0.67 | 3.48 ± 2.67 | 2.65 ± 2.28 | 2.32 ± 2.60 | 1.38 ± 2.05 |
| *Churning stomach* | 1.27 ± 0.80 | 1.30 ± 0.62 | 2.72 ± 2.29 | 2.20 ± 2.04 | 1.46 ± 2.22 | 0.91 ± 1.91 |
| *Fatigue* | 2.59 ± 1.83 | 2.87 ± 1.82 | 2.56 ± 1.79 | 2.61 ± 2.00 | -0.03 ± 2.26 | -0.28 ± 2.09 |
| *Feeling weak* | 1.63 ± 1.32 | 1.73 ± 1.05 | 3.68 ± 2.51 | 2.91 ± 2.09 | 2.02 ± 2.76 | 1.21 ± 1.82 |
| *Sweating* | 1.30 ± 0.76 | 1.64 ± 1.15 | 2.41 ± 2.01 | 2.29 ± 2.17 | 1.11 ± 1.90 | 0.63 ± 2.26 |
| *Muscle tension* | 1.98 ± 1.19 | 2.26 ± 1.41 | 4.46 ± 2.37 | 3.82 ± 2.34 | 2.48 ± 2.03 | 1.55 ± 2.11 |
| *Losing focus* | 1.86 ± 1.34 | 1.88 ± 1.42 | 3.45 ± 2.48 | 3.17 ± 2.33 | 1.58 ± 2.56 | 1.27 ± 2.19 |

#### 4.2 Change scores in self-reports across patients with stress-induced myocardial ischemia and without

Change scores for self-reported bodily sensations and emotions visualized across ischemic and non-ischemic patients are visualized in Figure S10. We sex-adjusted these scores by regressing out the sex-effect.

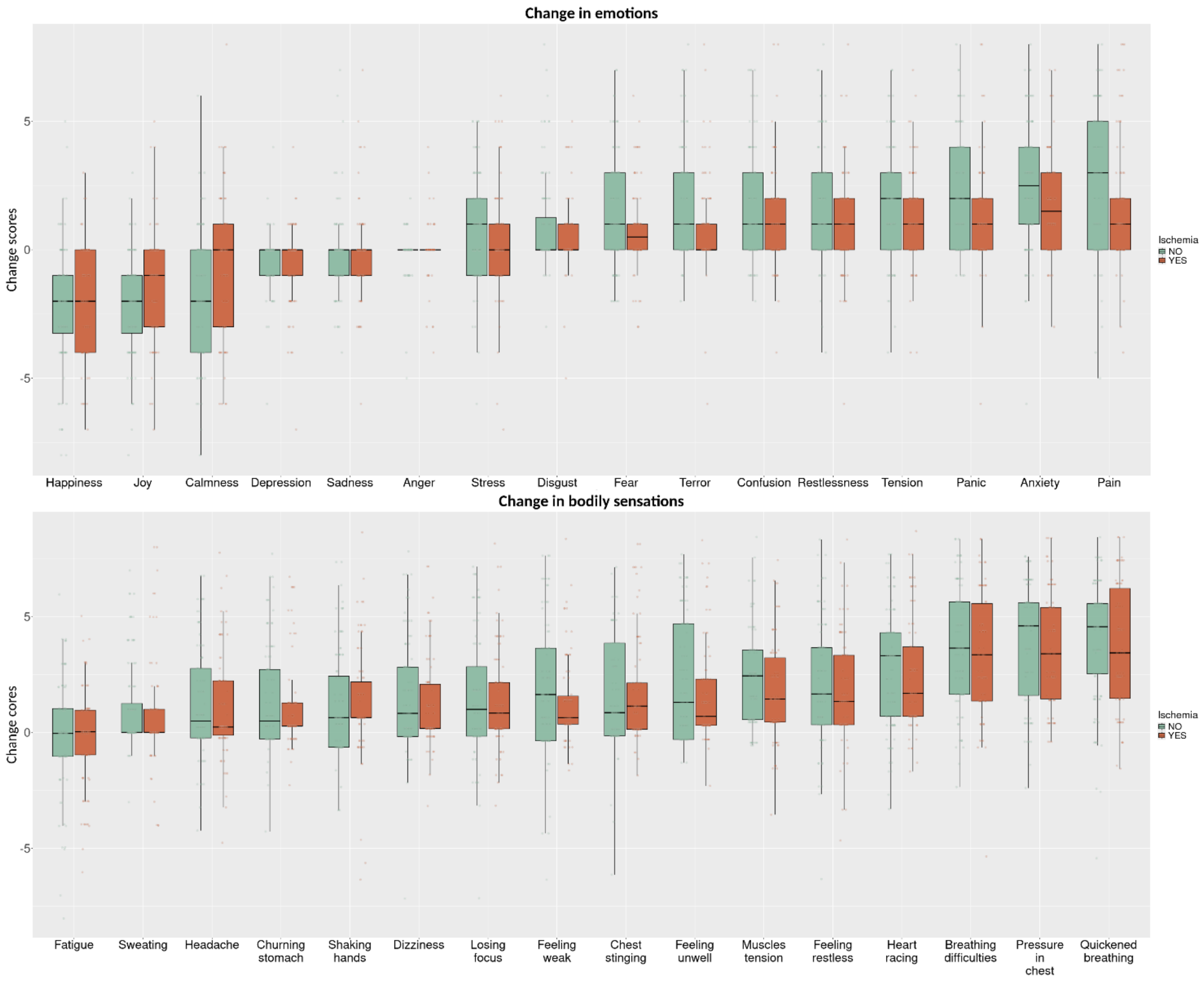


*Figure S10. Sex-adjusted change scores in emotional states (up) and bodily sensations (bottom) across ischemic and non-ischemic patients. Boxplot represents the interquartile range of the data. Whiskers show the smallest and the largest value within 1.5 times the interquartile range.*

### Appendix 5. Correlations between bodily sensations and emotions

We used Spearman's rank correlation to correlate changes in 16 emotional ratings and 16 bodily sensations with each other; matrices are visualized in **Figure S11**. The correlation matrix for associations between bodily sensations and emotions is shown in Figure 12.

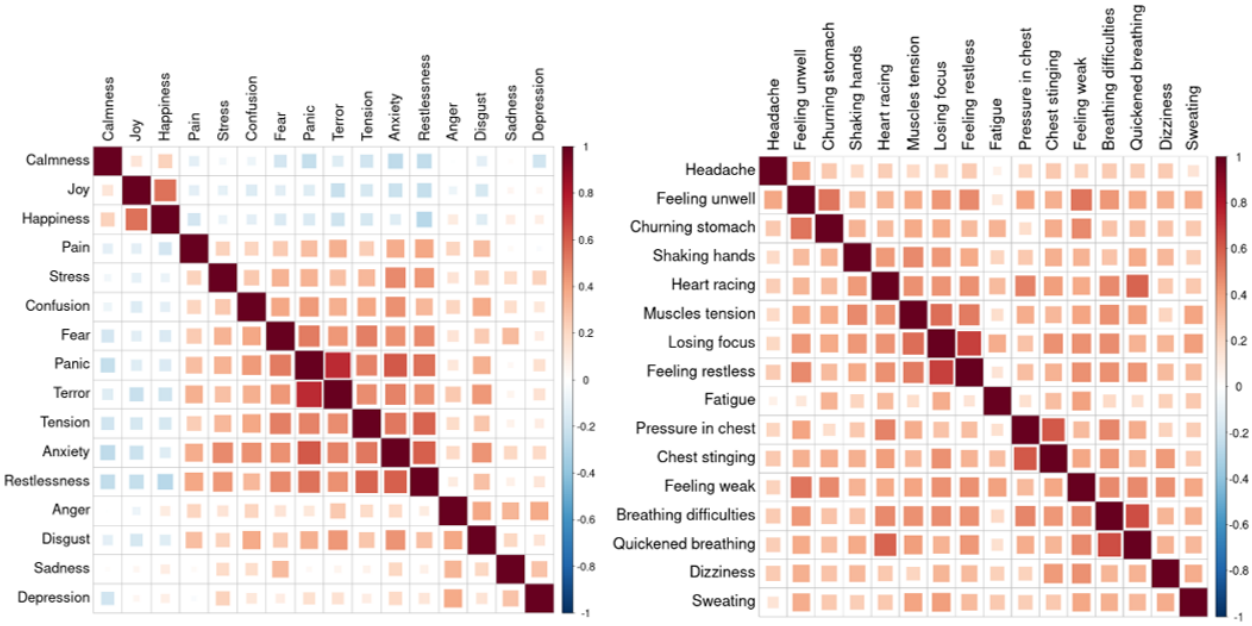


*Figure S11. Hierarchical Spearman correlation matrices between change in emotions (left) and change in bodily sensations (right) with each other.*

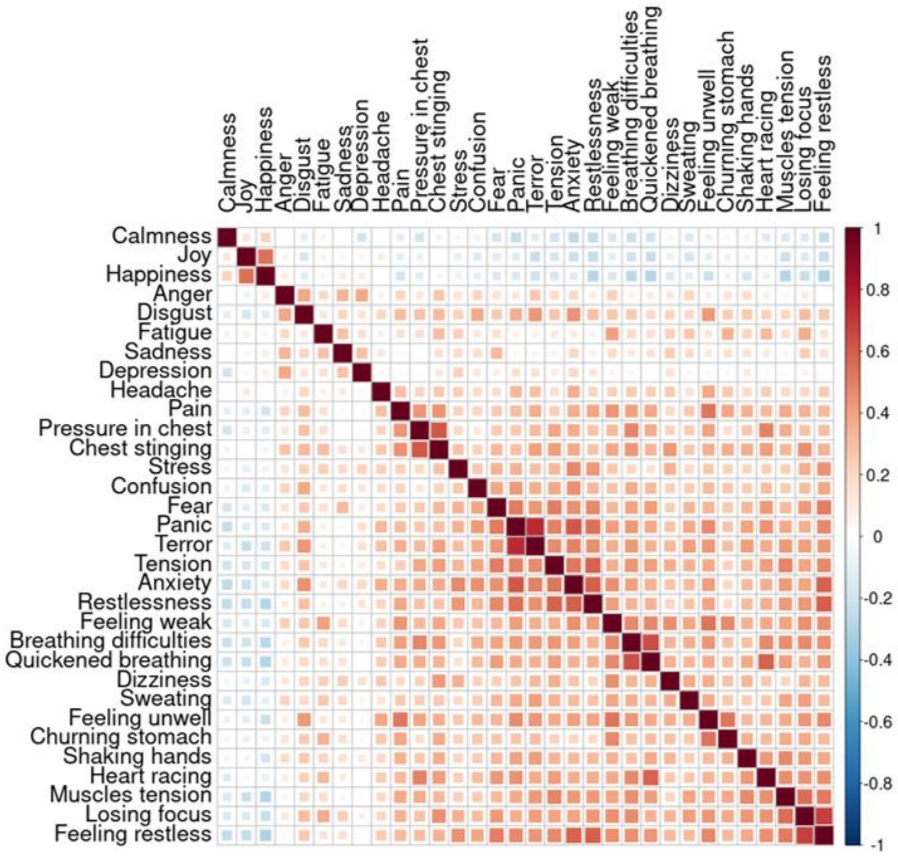


*Figure S12. Hierarchical Spearman correlation matrix between changes in 16 emotions and 16-bodily sensation.*

### Appendix 6. Hemodynamic effects

We investigated the effect of adenosine on hemodynamic measurements (heart rate, systolic and diastolic blood pressure), and how ischemic status and sex affect it (**Tables S5-S6**). We also examined the relationship between hemodynamic and psychological responses (**Table S7-11**).

#### 6.1. Linear modelling of the contribution of ischemic status and sex on hemodynamic parameters

We fit separate linear regression models for each hemodynamic outcome variable (HR, systolic blood pressure, and diastolic blood pressure) to estimate the fixed effects of sex and ischemic status. P-values were FDR-corrected.

*Table S5. Results of fitted linear models estimating the effect of ischemia and sex on the hemodynamic responses to adenosine. The table shows the beta estimate, STD error, t-value, p-value, and FDR corrected p-value. "Δ" refers to a change from the baseline value.*

| Hemodynamic variable | Predictor | Estimate | STD Error | t-value | p-value | pFDR |
|---|---|---|---|---|---|---|
| *peak HR* | Intercept | 99.14 | 1.69 | 58.53 | 8.36E-97 | 3.51E-95 |
| *peak HR* | SexM | -7.38 | 2.86 | -2.58 | 0.0109 | 0.020 |
| *peak HR* | IschemiaYES | -7.86 | 2.87 | -2.74 | 0.0070 | 0.014 |
| *Δ peak HR* | Intercept | 29.44 | 1.22 | 24.14 | 1.88E-50 | 9.87E-50 |
| *Δ peak HR* | SexM | 1.27 | 2.06 | 0.62 | 0.54 | 0.60 |
| *Δ peak HR* | IschemiaYES | -8.76 | 2.07 | -4.24 | 4.18E-05 | 0.00015 |
| *Δ HR 3-min* | Intercept | 21.36 | 1.27 | 16.76 | 1.33E-34 | 5.59E-34 |
| *Δ HR 3-min* | SexM | 1.84 | 2.15 | 0.86 | 0.39 | 0.46 |
| *Δ HR 3-min* | IschemiaYES | -9.10 | 2.16 | -4.21 | 4.66E-05 | 0.00015 |
| *Δ HR 6-min* | Intercept | 21.44 | 1.26 | 16.98 | 4.07E-35 | 1.90E-34 |
| *Δ HR 6-min* | SexM | -1.35 | 2.13 | -0.63 | 0.53 | 0.60 |
| *Δ HR 6-min* | IschemiaYES | -8.68 | 2.14 | -4.052 | 8.57E-05 | 0.00025 |
| *Δ DIA 3-min* | Intercept | -4.53 | 1.24 | -3.66 | 0.00037 | 0.00096 |
| *Δ DIA 3-min* | SexM | -3.02 | 2.09 | -1.45 | 0.15 | 0.20 |
| *Δ DIA 3-min* | IschemiaYES | 3.14 | 2.10 | 1.50 | 0.14 | 0.19 |
| *Δ SYS 3-min* | Intercept | -2.52 | 2.14 | -1.18 | 0.24 | 0.30 |
| *Δ SYS 3-min* | SexM | -0.13 | 3.60 | -0.036 | 0.97 | 0.97 |
| *Δ SYS 3-min* | IschemiaYES | -1.34 | 3.62 | -0.37 | 0.71 | 0.75 |
| *Δ DIA 6-min* | Intercept | -6.17 | 1.21 | -5.11 | 1.12E-06 | 4.26E-06 |
| *Δ DIA 6-min* | SexM | -3.61 | 2.04 | -1.77 | 0.079 | 0.12 |
| *Δ DIA 6-min* | IschemiaYES | 3.37 | 2.05 | 1.64 | 0.10 | 0.14 |
| *Δ SYS 6-min* | Intercept | -8.82 | 2.18 | -4.037 | 9.08E-05 | 0.00025 |
| *Δ SYS 6-min* | SexM | -1.76 | 3.68 | -0.48 | 0.63 | 0.68 |
| *Δ SYS 6-min* | IschemiaYES | 4.58 | 3.70 | 1.24 | 0.22 | 0.28 |
| *HR 3-min* | Intercept | 91.06 | 1.77 | 51.43 | 1.19E-89 | 2.50E-88 |
| *HR 3-min* | SexM | -6.80 | 2.99 | -2.28 | 0.024 | 0.043 |
| *HR 3-min* | IschemiaYES | -8.19 | 3.00 | -2.73 | 0.0072 | 0.014 |
| *HR 6-min* | Intercept | 91.15 | 1.78 | 51.26 | 1.82E-89 | 2.55E-88 |
| *HR 6-min* | SexM | -9.99 | 3.00 | -3.33 | 0.0011 | 0.0026 |
| *HR 6-min* | IschemiaYES | -7.77 | 3.01 | -2.58 | 0.011 | 0.020 |
| *DIA 3-min* | Intercept | 66.65 | 1.67 | 39.85 | 8.25E-76 | 4.95E-75 |
| *DIA 3-min* | SexM | -8.22 | 2.82 | -2.91 | 0.0042 | 0.0093 |
| *DIA 3-min* | IschemiaYES | 4.83 | 2.83 | 1.70 | 0.091 | 0.14 |
| *SYS 3-min* | Intercept | 132.20 | 2.96 | 44.66 | 6.18E-82 | 5.19E-81 |
| *SYS 3-min* | SexM | -8.27 | 4.99 | -1.66 | 0.10 | 0.14 |
| *SYS 3-min* | IschemiaYES | -1.04 | 5.02 | -0.21 | 0.84 | 0.86 |
| *DIA 6-min* | Intercept | 65.01 | 1.53 | 42.54 | 2.60E-79 | 1.82E-78 |
| *DIA 6-min* | SexM | -8.80 | 2.58 | -3.42 | 0.00085 | 0.0021 |

| | | | | | | |
|---|---|---|---|---|---|---|
| *DIA 6-min* | IschemiaYES | 5.059 | 2.59 | 1.95 | 0.053 | 0.086 |
| *SYS 6-min* | Intercept | 125.90 | 2.79 | 45.16 | 1.51E-82 | 1.59E-81 |
| *SYS 6-min* | SexM | -9.90 | 4.70 | -2.11 | 0.037 | 0.062 |
| *SYS 6-min* | IschemiaYES | 4.89 | 4.73 | 1.035 | 0.30 | 0.36 |

#### 6.2. Linear mixed effects model

We used linear mixed-effects models to investigate the effects of time point, sex, and ischemic status on hemodynamic measures. For each outcome, fixed effects included time (within-subject factor), sex, ischemic status, and their interactions (time × sex and time × ischemic status). A random intercept for participants was included to account for repeated measures. P-values were FDR adjusted.

*Table S6. Results of linear mixed effects model.*

| Hemodynamic variable | Predictor | Estimate | STD Error | DF | t-value | p-value | pFDR |
|---|---|---|---|---|---|---|---|
| *Heart rate* | Intercept | 69.70 | 1.67 | 185.7 | 41.74 | 2.65E-96 | 2.38E-95 |
| *Heart rate* | Time3min | 21.36 | 1.16 | 266 | 18.38 | 3.03E-49 | 1.64E-48 |
| *Heart rate* | Time6min | 21.44 | 1.16 | 266 | 18.45 | 1.65E-49 | 1.12E-48 |
| *Heart rate* | sexM | -8.64 | 2.82 | 185.7 | -3.069 | 0.0025 | 0.0061 |
| *Heart rate* | IschemiaYES | 0.90 | 2.83 | 185.7 | 0.32 | 0.75 | 0.81 |
| *Heart rate* | 3min:sexM | 1.84 | 1.96 | 266 | 0.94 | 0.35 | 0.47 |
| *Heart rate* | 6min:sexM | -1.35 | 1.96 | 266 | -0.69 | 0.49 | 0.63 |
| *Heart rate* | 3min:IschemiaYES | -9.096 | 1.97 | 266 | -4.62 | 6.12E-06 | 2.17E-05 |
| *Heart rate* | 6min:IschemiaYES | -8.68 | 1.97 | 266 | -4.40 | 1.55E-05 | 4.66E-05 |
| *Systolic pressure* | Intercept | 134.71 | 2.78 | 184.3 | 48.41 | 9.83E-107 | 2.65E-105 |
| *Systolic pressure* | Time3min | -2.52 | 1.92 | 266 | -1.31 | 0.19 | 0.27 |
| *Systolic pressure* | Time6min | -8.82 | 1.92 | 266 | -4.60 | 6.44E-06 | 2.17E-05 |
| *Systolic pressure* | sexM | -8.15 | 4.69 | 184.3 | -1.74 | 0.084 | 0.15 |
| *Systolic pressure* | IschemiaYES | 0.31 | 4.72 | 184.3 | 0.065 | 0.95 | 0.97 |
| *Systolic pressure* | 3min:sexM | -0.13 | 3.23 | 266 | -0.040 | 0.97 | 0.97 |
| *Systolic pressure* | 6minsexM | -1.76 | 3.23 | 266 | -0.54 | 0.59 | 0.69 |
| *Systolic pressure* | 3min:IschemiaYES | -1.34 | 3.25 | 266 | -0.41 | 0.68 | 0.76 |
| *Systolic pressure* | 6min:IschemiaYES | 4.58 | 3.25 | 266 | 1.41 | 0.16 | 0.24 |
| *Diastolic pressure* | Intercept | 71.18 | 1.57 | 184.7 | 45.47 | 3.17E-102 | 4.28E-101 |
| *Diastolic pressure* | Time3min | -4.53 | 1.081 | 266 | -4.19 | 3.79E-05 | 0.000102 |
| *Diastolic pressure* | Time6min | -6.17 | 1.081 | 266 | -5.71 | 2.95E-08 | 1.33E-07 |
| *Diastolic pressure* | sexM | -5.20 | 2.64 | 184.7 | -1.97 | 0.051 | 0.11 |
| *Diastolic pressure* | IschemiaYES | 1.69 | 2.65 | 184.7 | 0.64 | 0.53 | 0.65 |
| *Diastolic pressure* | 3min:sexM | -3.023 | 1.82 | 266 | -1.66 | 0.098 | 0.16 |
| *Diastolic pressure* | 6min:sexM | -3.61 | 1.82 | 266 | -1.98 | 0.049 | 0.11 |
| *Diastolic pressure* | 3min:IschemiaYES | 3.14 | 1.83 | 266 | 1.71 | 0.088 | 0.15 |
| *Diastolic pressure* | 6min:IschemiaYES | 3.37 | 1.83 | 266 | 1.84 | 0.067 | 0.13 |

#### 6.3 Time-series of hemodynamic measures during adenosine stress scan

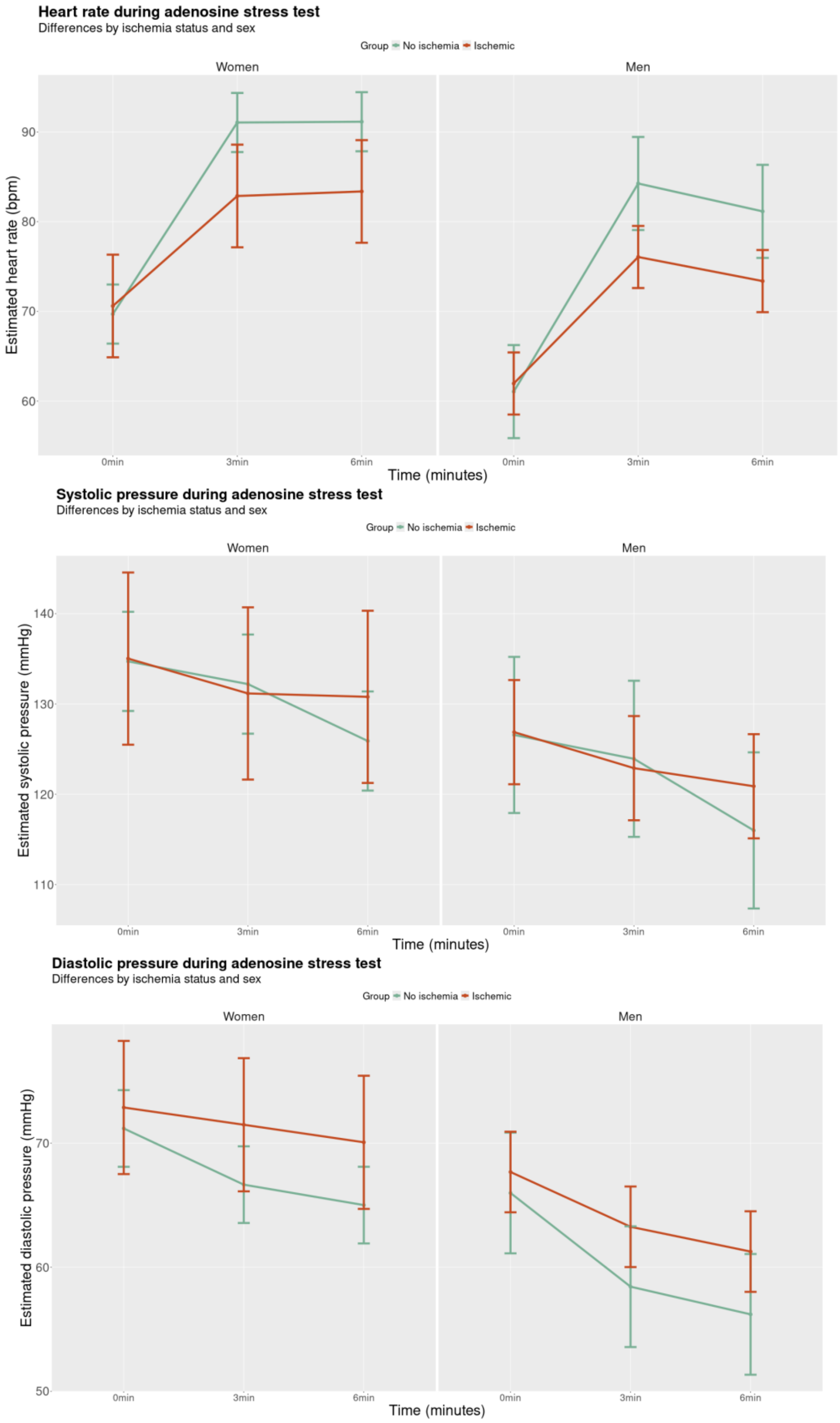


*Figure S13. Heart rate (top), systolic (middle), and diastolic blood pressure (bottom) at rest (0 min) and 3- and 6-minutes into adenosine stress testing, across sexes and ischemic status, separately for women (left) and men (right).*

In Figure S13, we visualized the time series of hemodynamic changes across the adenosine cardiac stress.

#### 6.4. Effects of cardiovascular parameters on the change in emotions and bodily sensations

We investigated the relationship between hemodynamic and psychological responses to adenosine. For that, we fit a separate linear regression model for each emotion and bodily sensation variable (response score). We investigate the effects on the peak HR response (rescaled by dividing by 10), and HR, systolic, and diastolic pressure values at three minutes into stress testing.

##### The effects of rescaled peak HR response

*Table S7. Effects of rescaled peak HR on reported changes in emotion and bodily sensation score. The table contains beta estimates, confidence intervals, p-values, and FDR-corrected p-values.*

| Psychological item | Beta estimate | Confidence interval | p-value | pFDR |
|---|---|---|---|---|
| *Quickened breathing* | 0.62 | [0.29, 0.95] | 0.00029 | 0.0036 |
| *Breathing difficulties* | 0.57 | [0.26, 0.88] | 0.00034 | 0.0036 |
| *Muscle tension* | 0.55 | [0.30, 0.80] | 3.15E-05 | 0.0010 |
| *Heart racing* | 0.47 | [0.18, 0.76] | 0.0015 | 0.0081 |
| *Anxiety* | 0.42 | [0.15, 0.69] | 0.0025 | 0.0089 |
| *Hands shaking* | 0.41 | [0.16, 0.67] | 0.0015 | 0.0081 |
| *Feeling restless* | 0.40 | [0.095, 0.71] | 0.011 | 0.024 |
| *Terror* | 0.38 | [0.14, 0.62] | 0.0023 | 0.0089 |
| *Sweating* | 0.37 | [0.11, 0.63] | 0.0057 | 0.016 |
| *Fear* | 0.36 | [0.13, 0.60] | 0.0022 | 0.0089 |
| *Confusion* | 0.35 | [0.10, 0.60] | 0.0058 | 0.016 |
| *Panic* | 0.34 | [0.083, 0.60] | 0.010 | 0.024 |
| *Sadness* | 0.33 | [0.14, 0.53] | 0.00094 | 0.0076 |
| *Tension* | 0.30 | [0.040, 0.56] | 0.024 | 0.051 |
| *Feeling restless* | 0.29 | [0.035, 0.55] | 0.026 | 0.053 |
| *Feeling unwell* | 0.28 | [-0.009, 0.57] | 0.057 | 0.11 |
| *Losing focus* | 0.25 | [-0.063, 0.55] | 0.12 | 0.18 |
| *Churning stomach* | 0.23 | [-0.017, 0.48] | 0.067 | 0.11 |
| *Chest stinging* | 0.22 | [-0.091, 0.52] | 0.17 | 0.24 |
| *Feeling of pain* | 0.21 | [-0.12, 0.54] | 0.21 | 0.27 |
| *Anger* | 0.19 | [0.064, 0.31] | 0.0033 | 0.011 |
| *Feeling weak* | 0.18 | [-0.12, 0.48] | 0.24 | 0.29 |
| *Pressure in chest* | 0.18 | [-0.12, 0.48] | 0.23 | 0.29 |
| *Disgust* | 0.18 | [-0.017, 0.39] | 0.072 | 0.12 |
| *Headache* | 0.16 | [-0.12, 0.43] | 0.26 | 0.31 |
| *Stress* | 0.11 | [-0.16, 0.38] | 0.40 | 0.44 |
| *Depression* | 0.059 | [-0.11, 0.23] | 0.50 | 0.53 |
| *Dizziness* | -0.0039 | [-0.27, 0.26] | 0.98 | 0.98 |
| *Calmness* | -0.0079 | [-0.32, 0.31] | 0.96 | 0.98 |
| *Fatigue* | -0.12 | [-0.40, 0.16] | 0.39 | 0.44 |
| *Joy* | -0.17 | [-0.43, 0.092] | 0.20 | 0.27 |
| *Happiness* | -0.26 | [-0.54, 0.016] | 0.065 | 0.11 |

#### Effects of HR in three minutes.

*Table S8. Effects of HR at three minutes into stress testing on reported changes in emotion and bodily sensation score. The table contains beta estimates, confidence intervals, p-values, and FDR-corrected p-values.*

| Psychological item | Beta estimate | Confidence interval | p-value | pFDR |
|---|---|---|---|---|
| *Quickened breathing* | 0.60 | [0.29, 0.92] | 0.00024 | 0.00378 |
| *Muscle tension* | 0.49 | [0.25, 0.73] | 0.00011 | 0.00355 |
| *Breathing difficulties* | 0.46 | [0.17, 0.76] | 0.0025 | 0.0112 |
| *Heart racing* | 0.46 | [0.18, 0.73] | 0.0014 | 0.00911 |
| *Sweating* | 0.44 | [0.19, 0.69] | 0.00059 | 0.00472 |
| *Fear* | 0.36 | [0.14, 0.58] | 0.0017 | 0.00911 |
| *Confusion* | 0.34 | [0.10, 0.58] | 0.0050 | 0.0177 |
| *Hands shaking* | 0.34 | [0.092, 0.58] | 0.0074 | 0.0236 |
| *Sadness* | 0.34 | [0.15, 0.52] | 0.00046 | 0.00472 |
| *Anxiety* | 0.29 | [0.021, 0.55] | 0.035 | 0.100 |
| *Feeling restless* | 0.29 | [-0.0061, 0.59] | 0.055 | 0.125 |
| *Panic* | 0.26 | [0.013, 0.51] | 0.040 | 0.100 |
| *Terror* | 0.25 | [0.010, 0.48] | 0.041 | 0.100 |
| *Feeling unwell* | 0.23 | [-0.049, 0.51] | 0.11 | 0.168 |
| *Churning stomach* | 0.22 | [-0.020, 0.44] | 0.072 | 0.145 |
| *Tension* | 0.22 | [-0.026, 0.47] | 0.078 | 0.148 |
| *Restlessness* | 0.21 | [-0.041, 0.46] | 0.10 | 0.168 |
| *Feeling weak* | 0.21 | [-0.080, 0.49] | 0.16 | 0.227 |
| *Disgust* | 0.18 | [-0.016, 0.37] | 0.073 | 0.145 |
| *Anger* | 0.18 | [0.057, 0.30] | 0.0041 | 0.0162 |
| *Losing focus* | 0.14 | [-0.16, 0.43] | 0.36 | 0.500 |
| *Headache* | 0.11 | [-0.16, 0.37] | 0.43 | 0.572 |
| *Feeling of pain* | 0.093 | [-0.22, 0.41] | 0.56 | 0.695 |
| *Stress* | 0.072 | [-0.19, 0.33] | 0.58 | 0.695 |
| *Pressure in chest* | 0.060 | [-0.23, 0.35] | 0.68 | 0.753 |
| *Dizziness* | 0.040 | [-0.22, 0.30] | 0.76 | 0.808 |
| *Depression* | 0.034 | [-0.13, 0.12] | 0.68 | 0.753 |
| *Chest stinging* | 0.032 | [-0.26, 0.33] | 0.83 | 0.842 |
| *Calmness* | 0.030 | [-0.27, 0.33] | 0.84 | 0.842 |
| *Happiness* | -0.22 | [-0.49, 0.046] | 0.10 | 0.168 |
| *Joy* | -0.18 | [-0.43, 0.069] | 0.16 | 0.227 |
| *Fatigue* | -0.074 | [-0.34, 0.19] | 0.59 | 0.695 |

#### Effects of systolic blood pressure at three minutes.

*Table S9. Effects of systolic pressure at three minutes into stress testing on reported changes in emotion and bodily sensation score. The table contains beta estimates, confidence intervals, p-values, and FDR-corrected p-values.*

| Psychological item | Beta estimate | Confidence interval | p-value | pFDR |
|---|---|---|---|---|
| *Anxiety* | 0.15 | [-0.012, 0.30 | 0.071 | 0.38 |
| *Fatigue* | 0.15 | [-0.0071, 0.31] | 0.061 | 0.38 |
| *Restlessness* | 0.15 | [-0.0026, 0.29] | 0.054 | 0.38 |
| *Muscle tension* | 0.14 | [-0.0083, 0.30] | 0.064 | 0.38 |
| *Panic* | 0.14 | [-0.0076, 0.29] | 0.063 | 0.38 |
| *Churning stomach* | 0.13 | [-0.010, 0.27] | 0.069 | 0.38 |
| *Tension* | 0.12 | [-0.033, 0.27] | 0.13 | 0.48 |
| *Feeling of pain* | 0.11 | [-0.075, 0.30] | 0.24 | 0.54 |
| *Quickened breathing* | 0.11 | [-0.086, 0.31] | 0.27 | 0.54 |
| *Stress* | 0.11 | [-0.041, 0.27] | 0.15 | 0.48 |
| *Terror* | 0.11 | [-0.034, 0.25] | 0.14 | 0.48 |
| *Sadness* | 0.10 | [-0.013, 0.22] | 0.083 | 0.38 |
| *Hands shaking* | 0.084 | [-0.066, 0.23] | 0.27 | 0.54 |
| *Breathing difficulties* | 0.083 | [-0.10, 0.27] | 0.37 | 0.66 |
| *Feeling weak* | 0.074 | [-0.099, 0.25] | 0.40 | 0.67 |
| *Fear* | 0.067 | [-0.070, 0.20] | 0.34 | 0.63 |
| *Confusion* | 0.058 | [-0.087, 0.20] | 0.43 | 0.69 |
| *Depression* | 0.058 | [-0.040, 0.16] | 0.24 | 0.54 |
| *Headache* | 0.056 | [-0.10, 0.21] | 0.48 | 0.74 |
| *Feeling restless* | 0.052 | [-0.13, 0.23] | 0.56 | 0.76 |
| *Chest stinging* | 0.049 | [-0.13, 0.23] | 0.59 | 0.76 |
| *Pressure in chest* | 0.043 | [-0.13, 0.22] | 0.63 | 0.76 |
| *Heart racing* | 0.042 | [-0.13, 0.21] | 0.63 | 0.76 |
| *Feeling unwell* | 0.037 | [-0.13, 0.21] | 0.66 | 0.76 |
| *Sweating* | 0.035 | [-0.12, 0.19] | 0.66 | 0.76 |
| *Anger* | 0.023 | [-0.050, 0.097] | 0.53 | 0.76 |
| *Calmness* | 0.0062 | [-0.17, 0.19] | 0.95 | 0.97 |
| *Dizziness* | 0.0040 | [-0.15, 0.16] | 0.96 | 0.97 |
| *Disgust* | 0.0030 | [-0.11, 0.12] | 0.96 | 0.97 |
| *Losing focus* | -0.0030 | [-0.18, 0.18] | 0.97 | 0.97 |
| *Joy* | -0.092 | [-0.24, 0.57] | 0.23 | 0.54 |
| *Happiness* | -0.095 | [-0.26, 0.66] | 0.25 | 0.54 |

#### Effects of diastolic pressure at three minutes

*Table S10. Effects of diastolic pressure at three minutes into stress testing on reported changes in emotion and bodily sensation score. The table contains beta estimates, confidence intervals, p-values, and FDR-corrected p-values.*

| Psychological item | Beta estimate | Confidence interval | P-value | pFDR |
|---|---|---|---|---|
| *Anxiety* | 0.47 | [0.20, 0.74] | 0.00087 | 0.028 |
| *Panic* | 0.38 | [0.12, 0.64] | 0.0050 | 0.079 |
| *Feeling restless* | 0.30 | [0.037, 0.56] | 0.026 | 0.22 |
| *Tension* | 0.29 | [0.032, 0.56] | 0.028 | 0.22 |
| *Terror* | 0.22 | [-0.023, 0.47] | 0.084 | 0.40 |
| *Sadness* | 0.20 | [-0.0039, 0.40] | 0.055 | 0.35 |
| *Breathing difficulties* | 0.19 | [-0.13, 0.512] | 0.24 | 0.78 |
| *Feeling restless* | 0.19 | [-0.12, 0.51] | 0.23 | 0.78 |
| *Muscle tension* | 0.17 | [-0.11, 0.44] | 0.23 | 0.78 |
| *Feeling of pain* | 0.14 | [-0.20, 0.47] | 0.42 | 0.87 |
| *Feeling unwell* | 0.13 | [-0.17, 0.43] | 0.39 | 0.87 |
| *Sweating* | 0.13 | [-0.15, 0.40] | 0.37 | 0.87 |
| *Feeling weak* | 0.12 | [-0.18, 0.43] | 0.43 | 0.87 |
| *Stress* | 0.12 | [-0.15, 0.40] | 0.38 | 0.87 |
| *Confusion* | 0.080 | [-0.18, 0.34] | 0.54 | 0.94 |
| *Shaking hands* | 0.080 | [-0.19, 0.34] | 0.56 | 0.94 |
| *Chest stinging* | 0.056 | [-0.26, 0.37] | 0.73 | 0.99 |
| *Fear* | 0.042 | [-0.20, 0.28] | 0.73 | 0.99 |
| *Heart racing* | 0.035 | [-0.27, 0.34] | 0.82 | 0.99 |
| *Quickened breathing* | 0.027 | [-0.32, 0.38] | 0.88 | 0.99 |
| *Pressure in chest* | 0.014 | [-0.29, 0.32] | 0.93 | 0.99 |
| *Disgust* | 0.0077 | [-0.20, 0.213] | 0.94 | 0.99 |
| *Depression* | 0.0065 | [-0.17, 0.18] | 0.94 | 0.99 |
| *Anger* | 0.0056 | [-0.12, 0.14] | 0.93 | 0.99 |
| *Churning stomach* | -0.0020 | [-0.25, 0.25] | 0.99 | 0.99 |
| *Losing focus* | -0.0066 | [-0.32, 0.31] | 0.97 | 0.99 |
| *Calmness* | -0.035 | [-0.35, 0.28] | 0.83 | 0.99 |
| *Dizziness* | -0.069 | [-0.34, 0.20] | 0.62 | 0.94 |
| *Fatigue* | -0.076 | [-0.36, 0.20] | 0.60 | 0.94 |
| *Headache* | -0.087 | [-0.37, 0.19] | 0.54 | 0.94 |
| *Happiness* | -0.11 | [-0.40, 0.17] | 0.43 | 0.87 |
| *Joy* | -0.23 | [-0.49, 0.033] | 0.087 | 0.40 |
